# Architectonic Principles of Polyproline II Helix Bundle Protein Domains

**DOI:** 10.1101/2023.11.25.568672

**Authors:** Cristian Segura Rodríguez, Douglas V. Laurents

## Abstract

Glycine rich polyproline II helix assemblies are an emerging class of natural domains found in several proteins with different functions and diverse origins. The distinct properties of these domains relative to those composed of α-helices and β-sheets could make glycine-rich polyproline II helix assemblies a useful building block for protein design. Whereas the high population of polyproline II conformers in disordered state ensembles could facilitate glycine-rich polyproline II helix folding, the architectonic bases of these structures are not well known. Here, we compare and analyze their structures to uncover common features. These protein domains are found to be highly tolerant of distinct flanking sequences. This speaks to the robustness of this fold and strongly suggests that glycine rich polyproline II assemblies could be grafted with other protein domains to engineer new structures and functions. These domains are also well packed with few or no cavities. Moreover, a significant trend towards antiparallel helix configuration is observed in all these domains and could provide stabilizing interactions among macrodipoles. Finally, extensive networks of Cα-H···O=C hydrogen bonds are detected in these domains. Despite their diverse evolutionary origins and activities, glycine-rich polyproline II helix assemblies share architectonic features which could help design novel proteins.

## Introduction

Protein structure comparison is a useful approach to uncover principles of protein architecture. By the mid-1970’s, a few dozen protein structures were known, which permitted Levitt and Chothia ^1^ to compare them via simplified topology/packing diagrams. Based on their results, they proposed an early structural classification scheme based on the content of protein secondary structure. They also found that when proteins are composed of just β secondary structure, the β-strands tend to be antiparallel and that in proteins of mixed α-helical and β-strand secondary structures, the α-helices tend to be positioned on the outside and β-strands tend to be parallel and on the inside. Moveover, certain preferred combinations of elements of secondary structure, such as two α-helices arranged in an antiparallel manner and two β-strands in parallel connected by an α-helix were discovered. Additionally, they uncovered that elements of protein secondary structure which are adjacent in the sequence are frequently in contact in the tertiary structure. Thanks to modern experiments in the protein design field, we now know that proximity of sequence and structures plays an important role in helping proteins fold quickly ^2^.

Further comparative studies identified the tendency of protein secondary structures to arrange into autonomously folding regions called domains in larger proteins and a general lack of knots^3^. Despite the relative success of protein comparative studies and new machine learning applications such as AlphaFold^4^ and RoseTTAFold^5^, they have largely focused on α-helices and β-sheet based protein structure and mostly ignore polyproline II helices ^6^.

Polyproline II helices are left-handed spirals with three residues per turn (**Figure 1A**). They are rather extended, as they advance 3.1 Å per residue ^7, 8^ compared to 1.5 Å per residue for α-helices and 3.0 – 3.8 Å per residue for β-strands^9^. The polyproline II (PPII) helix gets its name because this is the conformation adopted by peptides composed purely of all *trans* L-proline in aqueous solution ^10^. However, other residues can also adopt this conformation, which has backbone dihedral angles of about -75° for φ and +150° for ψ, which is near the basin for β-structure ( φ ≈ -135° , ψ ≈ +135) in the Ramachandran diagram. When every third proline residue is substituted by a glycine residue, three PPII helices can associate and H-bond, giving rise the abundant collagen triple helix structure (**Figure 1B**).

**Figure 1.**
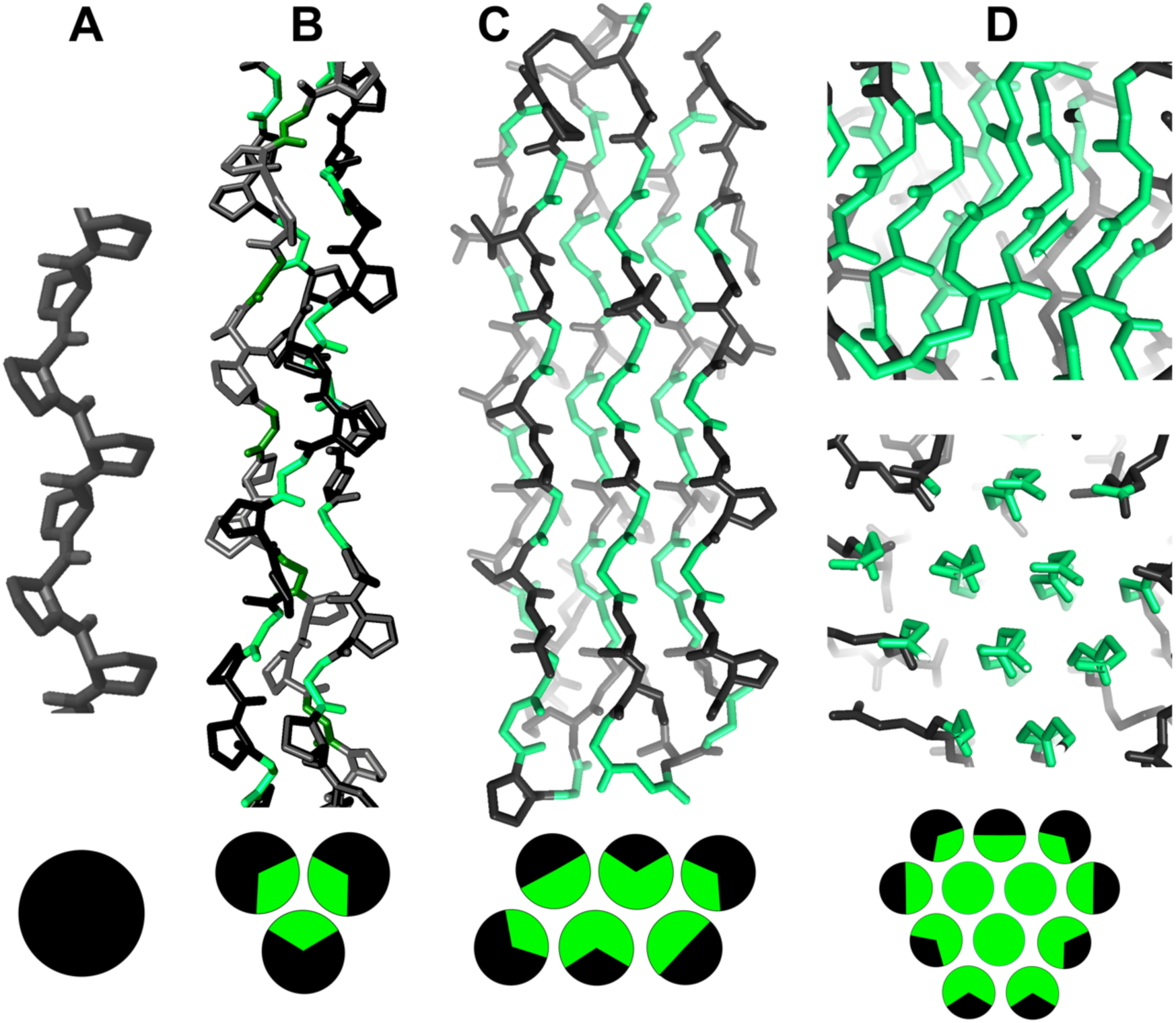
Proline and Glycine Rich Sequences Adopt Polyproline II Helices. **A.** Polyproline peptides adopt isolated polyproline II helices in aqueous solution. **B.** Substituting every third proline residue (black) for glycine (green) allows three chains to associate to form triple polyproline II helical structure of collagen. **C.** Swapping of additional prolines by glycine permits proteins with two layers of polyproline II helices to form, such as snow flea antifreeze proteins. **D.** Substitution of all proline residues (shown in black) by glycine (colored green) gives rise to a multilayered “honeycomb” of polyproline II helices, such as that seen in peptides composed of just glycine ^53^ and Human Anaplastic Lymphoma Kinase^64,65,75^.

Whereas collagen’s triple PPII helix structure is a standard of biochemistry textbooks, the presence of significant populations of PPII helical structure in short peptides and unfolded state ensembles is less well known. In the late 1960’s and early 1970’s, circular dichroism studies of polypeptides polyPro^11^, polyGlu and polyLys^12^ as well as collagen and globular proteins by Tiffany and Krimm provided the basis to propose specific spectral signatures for the disordered state and the polyproline II helix^13^. Moreover, they found that the PPII helix is adopted not only by poly(Pro) but also polyGlu and polyLys at pH values where their sidechains are charged^12^. The population of PPII helical conformations of these polypeptides, as well as some proteins, was found to increase upon cooling^14^ or by adding urea or guanidinium chloride^15^. These results were rationalized by arguing that the PPII helix conformation will reduce charge-charge repulsion in polyGlu and polyLys and effectively exposes the peptide backbone to solvent thus facilitating interaction with urea and guanidinium chloride. A few decades later, high populations of PPII helix were also detected in short peptide containing seven consecutive Ala residues by CD and NMR spectroscopies at low temperature; and underlined the importance of this conformation in unfolded state ensembles^16^. While controversial, these results were corroborated by 2D vibrational spectroscopy measurements of a trialanine peptide^17^. It was further confirmed by a very thorough investigation employing NMR spectroscopy and molecular dynamics simulations of a series of peptides containing three to seven consecutive alanine residues and established that these peptides at low temperature adopt a population of 90% PPII helix, 10% β-strand and essentially no α-helix^18^.

These remarkable findings led to research into the origin of the conformational stability of these PPII helices. Three effects have been proposed. First, Pappu and Rose and coworkers^19^ determined that contacts between non-sequential residues in peptide chains are significant and favor extended conformations, namely the β-strand and PPII helix regions of the Ramachandran diagram, over the α-helix. These results provided a new perspective on the Ramachandran diagram, which was developed based on steric clashes in dipeptides. Secondly, stabilizing *n*èπ* interactions can form between consecutive amide groups in the PPII helix but not in a β-strand conformation ^20^. Thirdly, PPII helical conformations are stabilized relative to other classes of secondary structure by backbone to water hydrogen bonding^21^ ^22^. While residues such as Val with bulky sidechains block somewhat this interaction and thus tend to favor β-strand-like extended conformations, others, like Gly or Ala whose side chains allow efficient water to backbone interactions, have particularly high propensities for polyproline II conformations^23,24,25^. Thus, it is now firmly established that the unfolded polypeptide chains contain sizeable populations of PPII helical conformations which impact their interactions and folding; see for example the excellent, thorough review by Schweitzer-Stenner ^26^.

Polyproline II helices play important roles in intrinsically disordered proteins (IDPs). For example, Titin is an enormous protein comprising a hundred folded domains linked by disordered proline-rich repeats which contain three short polyproline II helices^27^. The entropic stretching and recoil of these PPII helices and domains helps muscles spring back to their initial state following contraction^28^. As in short peptides, polyproline II helices in IDPs are most populated at lower temperatures ^29^. Detected on the residue level by NMR spectroscopy, partially populated PPII helices in Tau^30^, an amyloidogenic protein implicated in dementia, and CPEB, which forms a functional amyloid key for memory consolidation^31^, have been proposed to mediate physiologically important interactions with microtubules^30^ and profilin^32^, respectively, and to modulate these proteins’ formation of β-strand-rich amyloid structures. In the protein huntingtin, the pathologically relevant amyloidogenic polyglutamine segment is followed by a shorter stretch of polyproline. Several lines of evidence suggest that the polyproline stretch forms PPII helices which prevent amyloid formation by interacting with the polyglutamine segment through a combination of hydrophobic interactions and hydrogen bonding between Gln sidechain -NH_2_ and Pro backbone O=C moieties ^33^. PPII helices can also promote the formation of biomolecular condensates such as that formed by nephrin/Nck/N-WASP in the kidney’s glomerular filtration barrier^34^. As will be discussed below, glycine rich segments can also adopt polyproline II helical assemblies. Glycine rich segments are present in some proteins, such as FUS, and the possible role of these glycine rich segments forming networks of PPII helices or other assemblies which promote biomolecular condensate formation has been proposed^35,36,37^. Moreover, glycine residues in the GxxxG motif facilitate dimerization of α-helices in a large number of membrane proteins^38^, including the Aβ amyloid polypeptide ^39^. As in PPII helical protein domains rich in glycine, Cα-H··O=C interactions have been reported to form in GxxxG motifs in the Aβ amyloid polypeptide and impact pathology^40^. Moreover, Cα-H··O=C H-bonds formed by this motif can also affect the α-helix – α-helix packing angle and α-helical stability in model α-helices assembled in membrane lipids ^41^.

Polyproline II helices are also common in folded proteins. In addition to unfolded state ensembles and intrinsically disordered proteins, the “loop” segments connecting α-helices and β-sheets frequently adopt PPII helices as detected by X-ray crystallography studies^42^. Polyproline II helices in folded proteins can also be detected and quantified by far ultraviolet circular dichroism spectroscopy^43^. Certain residues, especially Ala, Gly, and Pro prefer to adopt PPII helical conformations in protein “loop” segments whereas the branched and aromatic residues tend to adopt β-strand-like conformations. Whereas water molecules forming bridging H-bonds between different protein H-bond donor and acceptor groups do not seem to be responsible for these preferences^44^, they have been attributed to an electrostatic screening model by which the side chain impacts the interaction between the backbone and the solvent ^23^. The same trends have been detected spectroscopically in short peptides^24^. These PPII helices are usually solvent-exposed and often mediate protein/protein, protein/nucleic acid and pathogen/host interactions ^8,45^, including that between the SARS-CoV-2 spike protein and the ACE2 receptor ^46^. In addition to these isolated PPII helices on the surface of globular proteins, several proteins, including the complement C1q^47^, SPP1 viral capsid protein ^48^ and the synaptic acetylcholineesterase^49^ have been found to contain collagen-like segments which serve as trimerization motifs.

Our improving understanding of PPII helices is facilitating their use in protein design. For example, the successful engineering of collagen-like triple helices into higher order hollow octadecameric assemblies through additional contacts on the outer face of some helices has been recently achieved^50^. Higher level structures should also be possible since peptides containing the sequence (Pro-Gly-Gly)_N_ adopts polyproline II helices which assemble into double-layered sheets with proline residues on the outside and glycine residues and an extensive hydrogen bonding network on the inside^51^ (**Figure 1C**). A similar structure is adopted by (Ala-Gly-Gly)_N_^52^. Remarkably, the substitution of all three prolines for glycine gives rise to the complete 3D network of PPII helices in which each PPII helix is surrounded by and forms hydrogen bonds to six others ^53^ (**Figure 1D**). The presence of these N-H··O=C ^53^ and Cα-H··O=C^54^ hydrogen bonds, was corroborated by infrared and Raman spectroscopies^55^. A closely related structure is adopted by (Ala-Gly-Gly-Gly)_N_ ^52^. Whether these chains have a preference for a parallel or antiparallel orientation remains an open question. Except for collagen and collagen-like triple helices with one third glycine, throughout the 1970’s, 1980’s and 1990’s the sequences and conformations described above with two thirds and even 100% glycine were not observed in natural proteins and seemed to be academic curiosities found only in synthetic peptides. This view changed in the last two decades as various natural protein domains and proteins have been found to be composed of several glycine-rich PPII helices arranged in a bundle. They include the antifreeze proteins from primitive arthropods ^56,57^ a universally conserved translation factor ^58, 59, 60^, enzymes capable of forming new carbon – carbon bonds to incorporate CO_2_ into organic compounds ^61,62^, the key tip domain in bacteriophage tail fibrils ^63^, the anaplastic lymphoma kinase ^64^ and the leukocyte tyrosine kinase^65^. Their names and some characteristics are listed in **Table 1**.

**Table 1.**
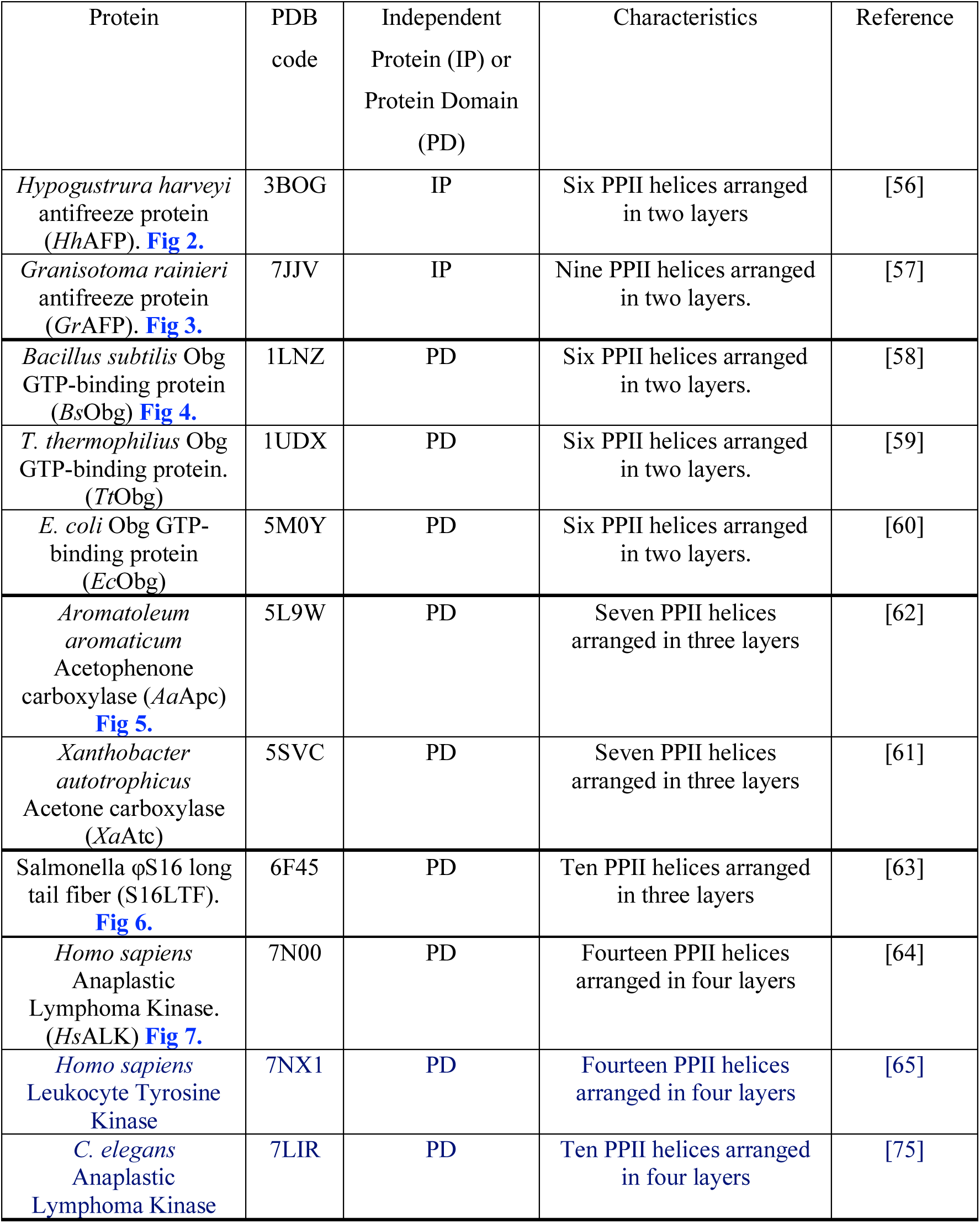

In particular, the antifreeze proteins have proven valuable as systems to investigate the bases of the conformational stability of glycine-rich PPII helical bundles and have revealed disulfide bonds, nonconventional Cα-H···O=C hydrogen bonds, and a lower than anticipated conformational entropy in the denatured state as factors favoring the folded structure ^66,35,67^. Nevertheless, some important questions remain regarding the roles of : 1) flanking sequences, which are known to be important for amyloid structures ^68^, 2) packing density, which is expected to impact the stabilizing contribution of van der Waals interactions, 3) the parallel versus antiparallel orientation of the helices^53^, which we hypothesize could be stabilizing or destabilizing due to macrodipole interactions and 4) unconventional Cα-H···O=C hydrogen bonds.

Thanks to the diverse evolutionary origin, function and complexity of the PPII helical protein domains reported recently, a comparative study of their structural features is now possible. Following the approach pioneered by Levitt & Chothia ^1^, and by characterizing helix orientation and the sequence composition of flanking segments, we aim to address the questions posed above to uncover architectural principles for PPII helical bundle domains. This new knowledge may help guide the design of novel proteins based on PPII helical assemblies.

## Methods

X-ray crystal structures of PPII helical bundle proteins and domains were examined and represented as 2D topological diagrams following the approach of Levitt & Chothia (1976) ^1^. We tabulated the residues in the turns connecting the PPII helices to uncover characteristics of flanking residues. For longer connecting sequences, the five residues immediately preceding and the N-terminus and the five residues immediately following the C-terminus of each PPII helix were tabulated. The proteins’ packing densities at buried positions in PPII helices were evaluated with the program Voronoia^69^; for comparison, the same analysis was also applied to α-helices and β-strands in some representative proteins previously studied by Liang and Dill^70^.

Glycine residues have two Hα and in NMR spectra the chemical shifts of these two hydrogens become distinct when a hydrogen bond is formed. In *Aa*AFP, there are 14 Gly residues whose two Hα show chemical shifts differing by 0.50 ppm or more. This number agrees well with the 15 Cα-H participating in H-bonds identified by Gates *et al.* (2017)^66^. These Gly residues have one Hα positioned to accept a hydrogen bond from a nearby carbonyl group. Considering that for these H-bonds, the <u>Cα-H··O</u>=C distance is 3.54 +/-0.2 (1σ) Å and the <u>Cα-H···O</u>=C bond angle is 157 +/-22° (1σ), as criteria to detect putative Cα-H···O=C H-bonds in other glycine-rich PPII helical bundle proteins, we used a distance limit of < 3.75 Å (3.54 Å + 1σ) and a bond angle > 135° (157° – 1σ). The search was carried out using the program PYMOL and each detected putative Cα-H··O=C H-bond was subsequently checked and curated manually. The structures analyzed here range from very high to medium resolution. In particular, two of the structures studied (3BOG & 7JJV) are at very high “atomic resolution” of 1.2 Å and 1.21 Å, one is high resolution (6F45 at 1.70 Å), two are at medium resolution: 1LNZ at 2.6 Å and 5L9W at 2.9 Å and the last structure (PDB: 7N00) was solved by cryoEM with a reported resolution of 2.27 Å. This is pertinent since the uncertainty in H-bond lengths is known to vary with structure resolution^71^. Using reported data,^70^ the H-bond uncertainties are estimated to be 0.12 Å for 3BOG and 7JJV, 0.15 Å for 6F45, 0.23 Å for 1LNZ, 0.26 Å for 5L9W and 0.18 Å for 7N00. Considering the lower resolution and higher bond length uncertainties in 1LNZ, 5L9W and 7N00, the number of hydrogen bonds detected here may be underestimated.

## Results and Discussion

Topological diagrams for all reported polyproline II helical bundle proteins are shown in **Figure 2-7** in order of increasing complexity alongside their sequences and 3D structures. The snow flea antifreeze proteins have the simplest structures, which consist of two layers of PPII helices connected by short loops (**Fig. 2 & 3**). The two disulfide bonds present in *Hh*APF contribute decisively for the protein’s conformational stability. The PPII helices alternate having their N- or C-termini up or down and this antiparallel arrangement would promote favorable electrostatic interactions between the helix macrodipoles. The N- and C-terminal PPII helices have only two other PPII helices as neighbors and are relatively poor in Gly residues.

**Figure 2.**
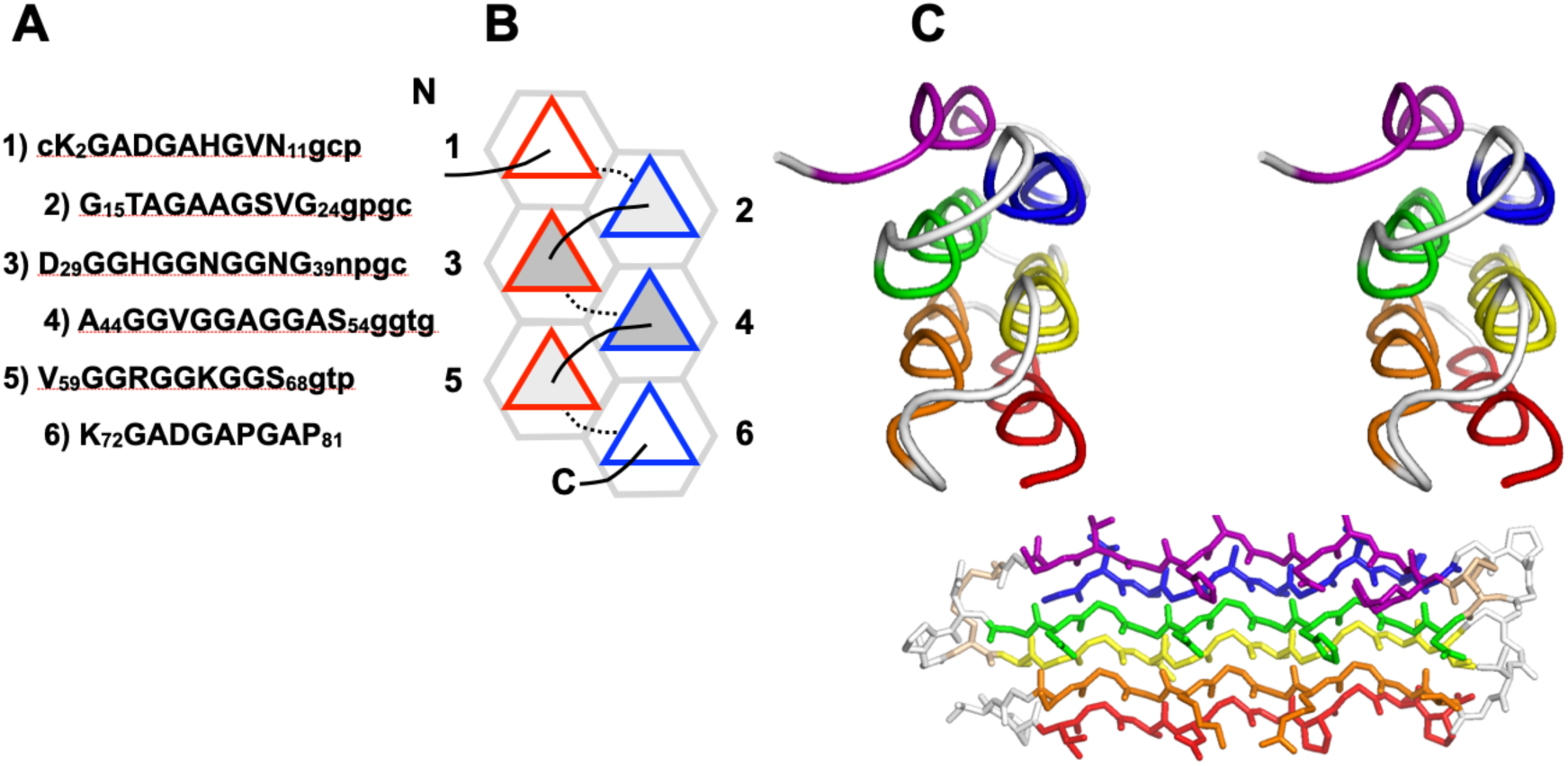
*Hypogustrura harveyi* antifreeze protein (*Hh*AFP). **A.** Primary structure of *Hf*AFP. Residues in PPII helices (which are numbered) are shown in capital letters, other residues in the connecting loops are written in lowercase. **B.** Schematic diagram representing the PPII helices as triangles. Connecting loops above the plane are shown as solid lines; those below the plane are shown as dotted lines. PPII helical N-termini pointing up are outlined in red; N-termini pointing down are outlined in blue. PPII helices with two, three and four neighboring helices are shaded white, light gray and medium gray, respectively. **C.** (*top*) Cross-eyed stereo ribbon diagram of the *Hh*AFP tertiary structure (PDB 3BOG) colored blue to red from the N- to the C-terminus, with connecting turns in gray, showing the six PPII helices end on (*top*) and along the long axis (*bottom*).

**Figure 3.**
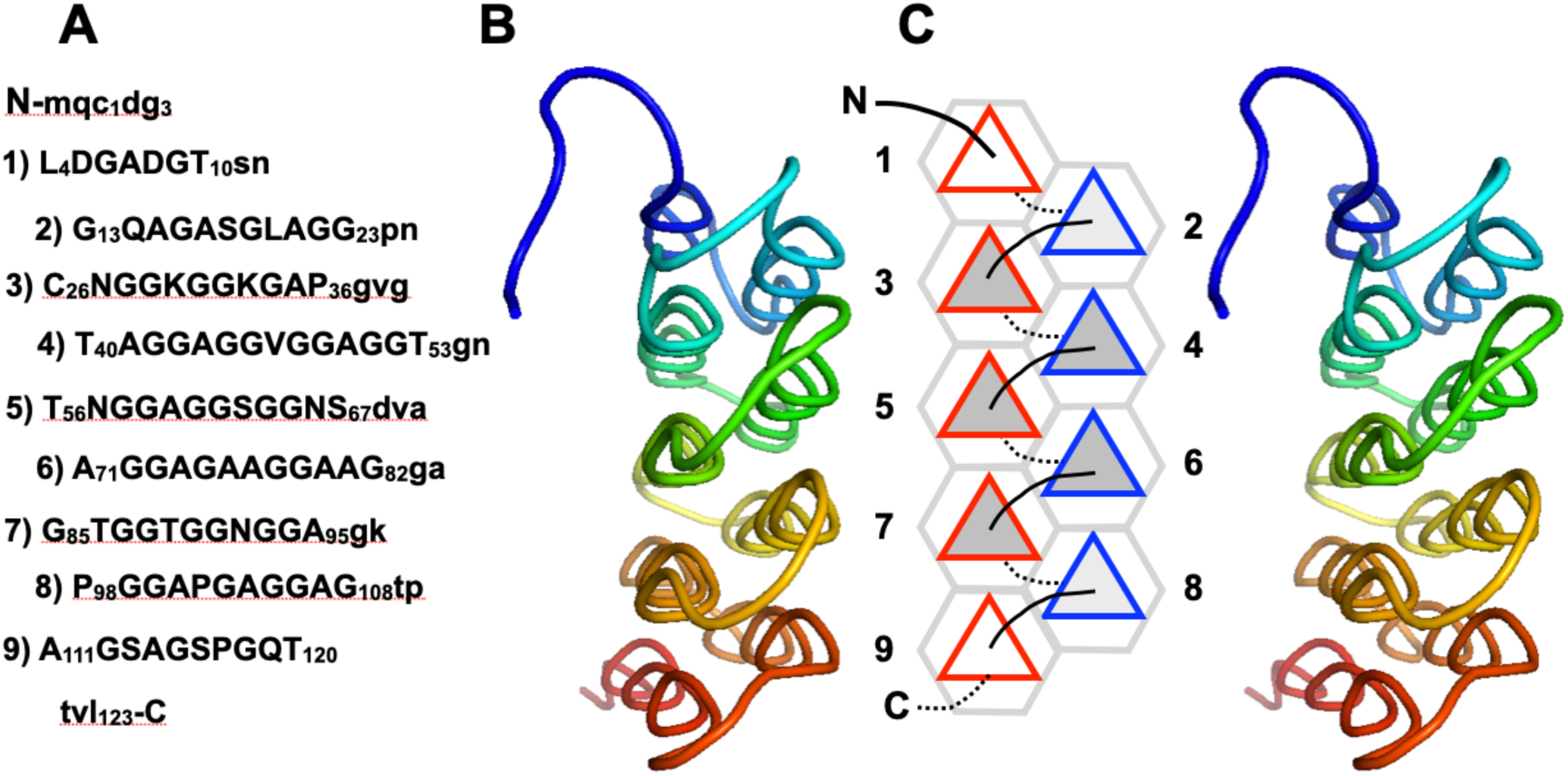
*Granisotoma rainieri* antifreeze protein. **A.** Primary structure of *Gr*AFP. Residues in PPII helices (which are numbered) are shown in capital letters, others are written in lowercase. **B.** Cross-eyed stereo ribbon diagram of the *Gr*AFP tertiary structure (PDB 7JJV) colored blue to red from the N- to the C-terminus, showing the nine PPII helices end on. **C.** Schematic diagram representing the PPII helices as triangles. Connecting loops above the plane are shown as solid lines; those below the plane are shown as dotted lines. PPII helical N-termini pointing up are outlined in red; N-termini pointing down are outlined in blue. PPII helices which two, three and four neighboring helices are shaded white, light gray and medium gray, respectively.

**Figure 4.**
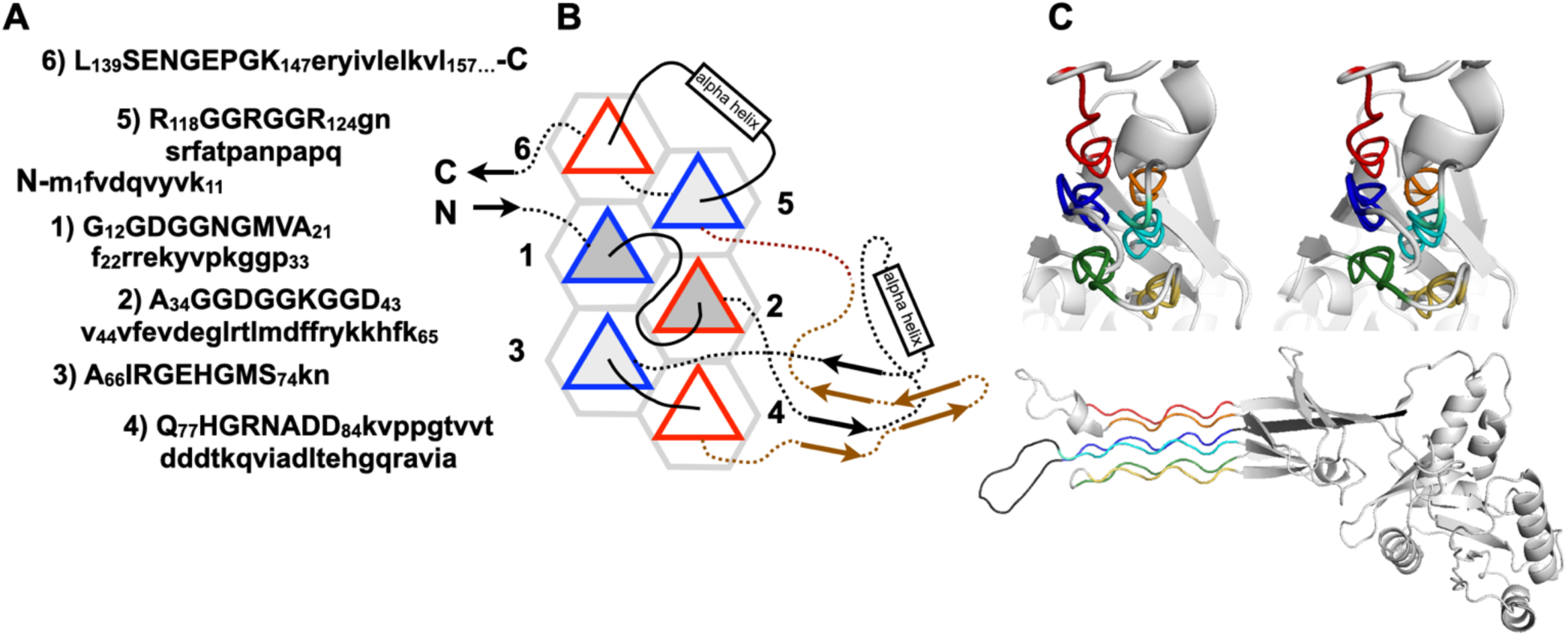
*Bacillus subtilis* Obg GTP-binding protein (*Bs*Obg) **A.** Primary structure of *Bs*Obg. Residues in PPII helices (which are numbered) are shown in capital letters, others are written in lowercase. **B.** Schematic diagram representing the PPII helices as triangles. Connecting loops above the plane are shown as solid lines; those below the plane are shown as dotted lines. For clarity, the connections between PPII helices 4 and 5 are colored brown. PPII helical N-termini pointing up are outlined in red; N-termini pointing down are outlined in blue. PPII helices which two, three and four neighboring helices are shaded white, light gray and medium gray, respectively. **C.** Cross-eyed stereo ribbon diagram of the *Bs*Obg tertiary structure (PDB 1LNZ) colored blue to red from the N-to the C-terminus, showing the six PPII helices end on (top). The six PPII helices together with the rest of the protein (light gray) are shown in the lower panel.

**Figure 5.**
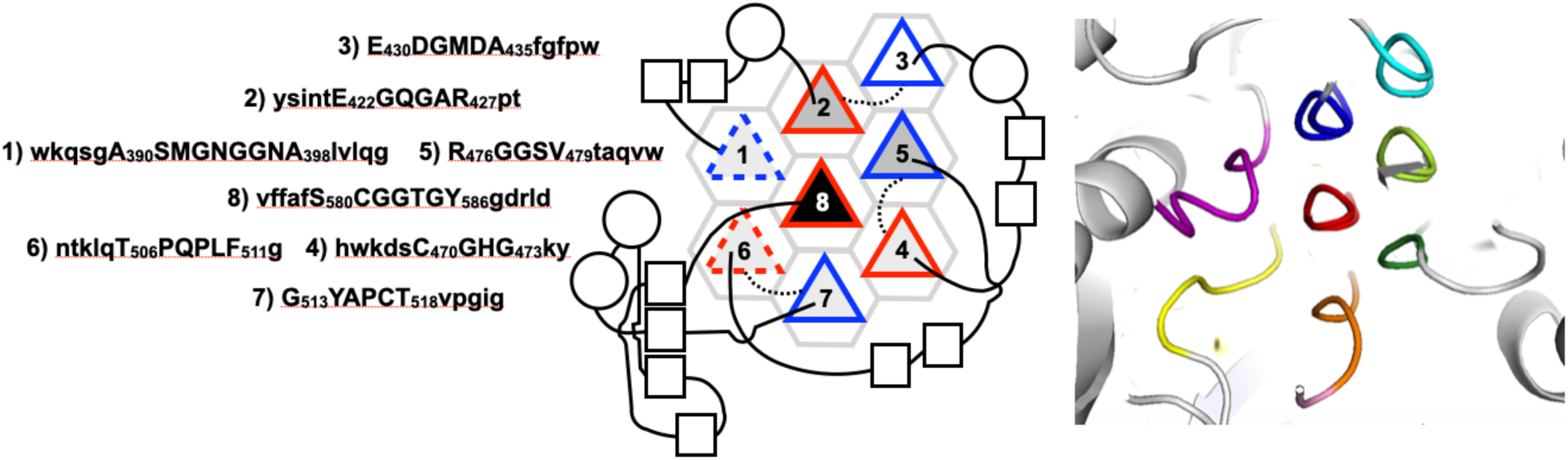
*Aromatoleum aromaticum* Acetophenone Carboxylase (*Aa*APC) **A.** The primary structure of *Aa*APC PPII helices is shown, with the PPII helices numbered. Residues forming the PPII helices are written in capital letters; those of flanking sequence are shown in lower case. **B.** Schematic diagram representing the PPII helices as triangles. Connecting loops above the plane are shown as solid lines; those below the plane are shown as dotted lines. PPII helical N-termini pointing up are outlined in red; N-termini pointing down are outlined in blue. The dashed outline of PPII helices 1 and 6 refers to their more deviant geometry. PPII helices with two, three, four and six neighboring helices are shaded white, light gray, medium gray and black, respectively. α-helices and β-strands of the connecting segments are represented as circles and squares, respectively. Squares which are aligned and close together represent β-strands hydrogen bonded in β-sheets. **C.** The *Aa*APC tertiary structure (PDB 5L9W) with PPII helices colored purple, blue, cyan, dark green, lime green, yellow, orange and red from the N- to the C-terminus, showing the eight PPII helices end on.

**Figure 6.**
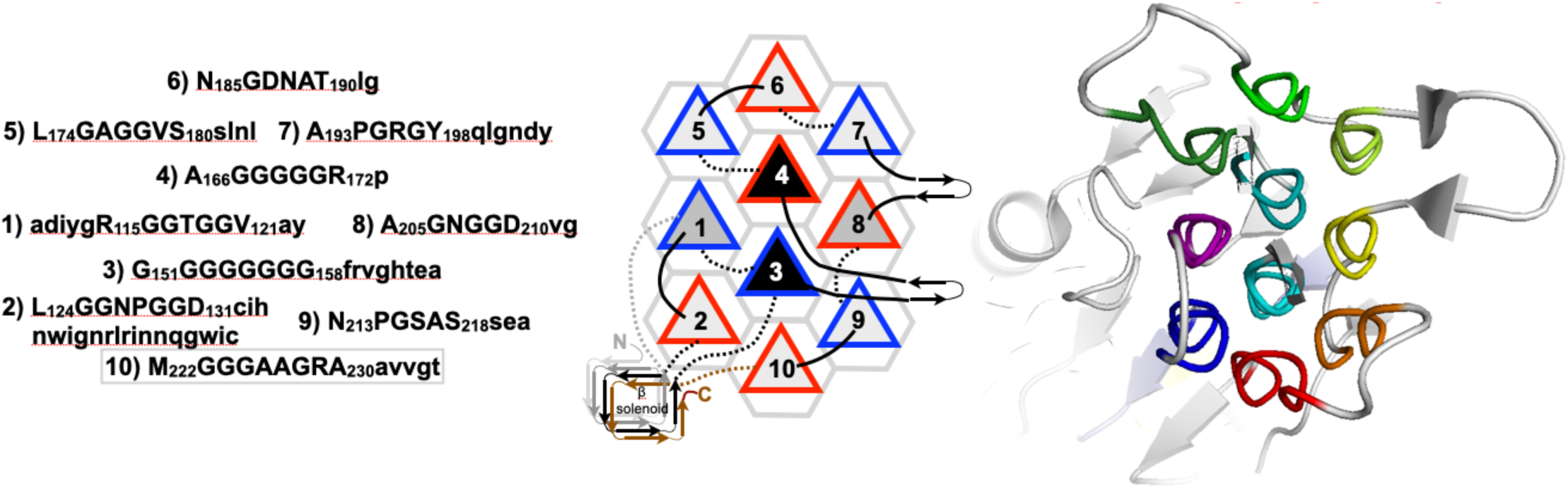
*Salmonella* φS16 Long Tail Fiber gp38 Domain (S16LTF). **A.** The primary structure of S16LFT gp38 domain PPII helices and flanking segments is shown, which PPII helices labeled. Residues forming the PPII helices are written in capital letters, those forming the connecting segments are written in lowercase. **B.** Schematic diagram representing the PPII helices as triangles. Connecting loops above the plane are shown as solid lines; those below the plane are shown as dotted lines. PPII helical N-termini pointing up are outlined in red; N-termini pointing down are outlined in blue. PPII helices with three, four and six neighboring helices are shaded light gray, medium gray and black respectively. PPII helix 1 is preceded by two turns of a β-solenoid (gray), another layer of this β-solenoid is contributed by the segment linking PPII helices 2 and 3 (black) and a final layer is formed by the segment following PPII helix 10 (brown). **C.** The S16LFT tertiary structure (PDB 6F45) colored purple to red from the N- to the C-terminus, showing the ten PPII helices end on.

**Figure 7.**
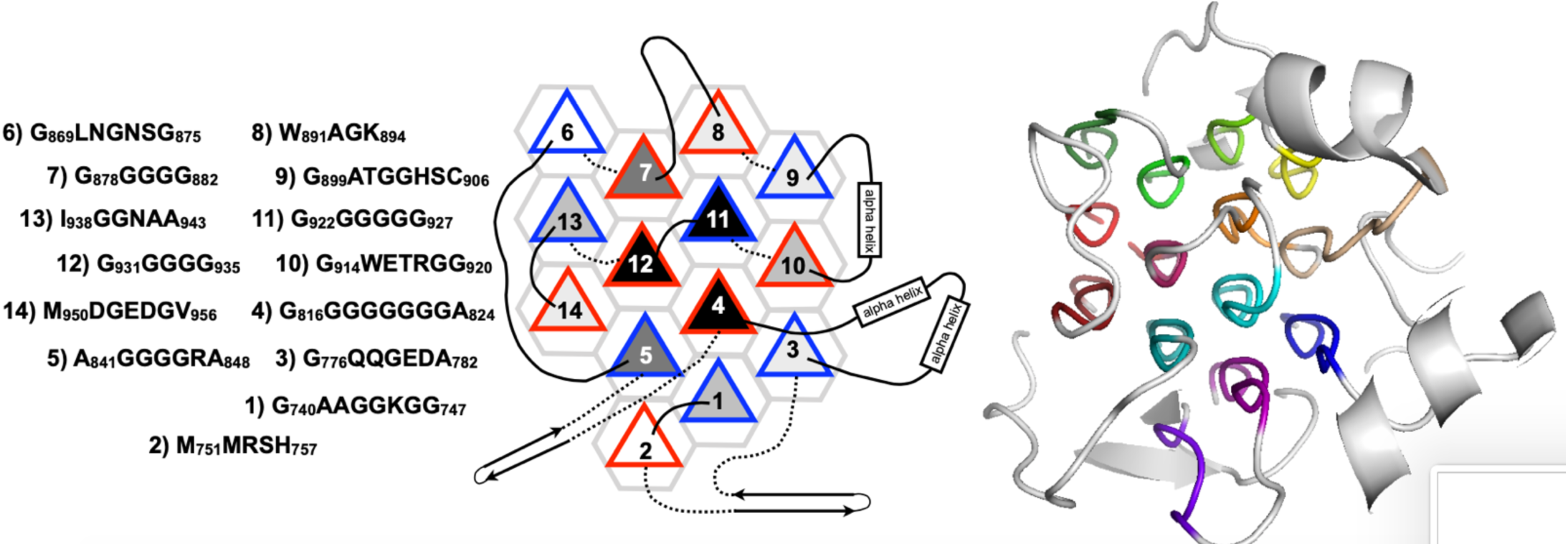
*Homo sapiens* Anaplastic Lymphoma Kinase (*Hs*ALK) **A.** Primary structure of the fourteen PPII helices in *Hs*ALK. Helices are numbered according to the order along the sequence. **B.** Schematic diagram representing the PPII helices as triangles. Connecting loops above the plane are shown as solid lines; those below the plane are shown as dotted lines. For clarity, the sequence of the connecting segments are not shown. PPII helical N-termini pointing up are outlined in red; N-termini pointing down are outlined in blue. PPII helices which two, three and four, five and six neighboring helices are shaded white, light gray, medium gray, dark gray and black respectively. PPII helices with 5 or 6 neighbors are labeled in white. **C.** The tertiary structure of the PPII helical bundle domain of *Hs*ALK (PDB: 7N00) is shown end-on with helices colored in a rainbow spectrum from purple (helix 1) to dark red (helix 14).

Like *Hh*AFP, *Bs*Obg is also composed of six PPII helices (**Fig. 4**) arrange in two layers. However, these PPII helices are connected by segments of varying length and complexity; some even include α-helices and β-strands. The PPII helical domain of *Bs*Obg is not connected by disulfide bonds and are also arranged to promote favorable PPII macrodipole interactions. Like the snow flea AFPs, the central PPII helices, which have four bordering PPII helices, tend to contain more Gly residues.

Acetophenone carboxylase and acetone carboxylase are enzymes which incorporate CO_2_ in the form of bicarbonate into organic compounds through the formation of new carbon-carbon bonds:

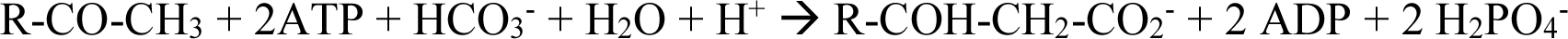

Where “R” is a methyl or phenyl group in the case of acetone carboxylase or acetophenone carboxylase, respectively. For acetone carboxylase, one ATP is converted into one AMP and two P_i_. These enzymes contain a PPII domain consisting of eight rather short PPII helices ^61,62^. Remarkably, one PPII helix is surrounded by six others and has a high glycine content and the whole PPII helical bundle domain is buried and surrounded by other enzyme domains^61,62^. These structures show that PPII helical bundle domains need not be long or solvent exposed, which would afford water molecules as H-bonding partners, to be stable.

The bacteriophage S16, which infects Salmonella bacteria, has a long tail fiber composed of several domains. The last domain (gp38), right at the fiber’s tip, is composed of a complex glycine-rich PPII helical bundle ^63^. Composed of ten PPII helices, this domain included two PPII helices which are completely surrounded by other PPII helices and are composed completely of glycine (**Figure 6**). In fact, for a given PPII helix, its glycine content in all these proteins increases with the number neighboring PPII helices (**Figure 8A**). The phage gp38 domain’s amino acid sequence is well conserved across T-even bacteriophages (except T4). It serves as a stable platform for five surface loops which are highly variable in length and sequence. These loops recognize outer membrane proteins on the surface of prey bacteria which is the key initial event in infection. Residue substitutions in these loops have been shown to alter prey specificity. Amyloid stability and polymorphism is frequently impacted by the sequence of neighboring segments. By contrast, the PPII helical bundle of the phage gp38 domain remains stable despite the variation of the surface loops’ sequence and this speaks to its conformational stability and the robustness of the fold.

**Figure 8.**
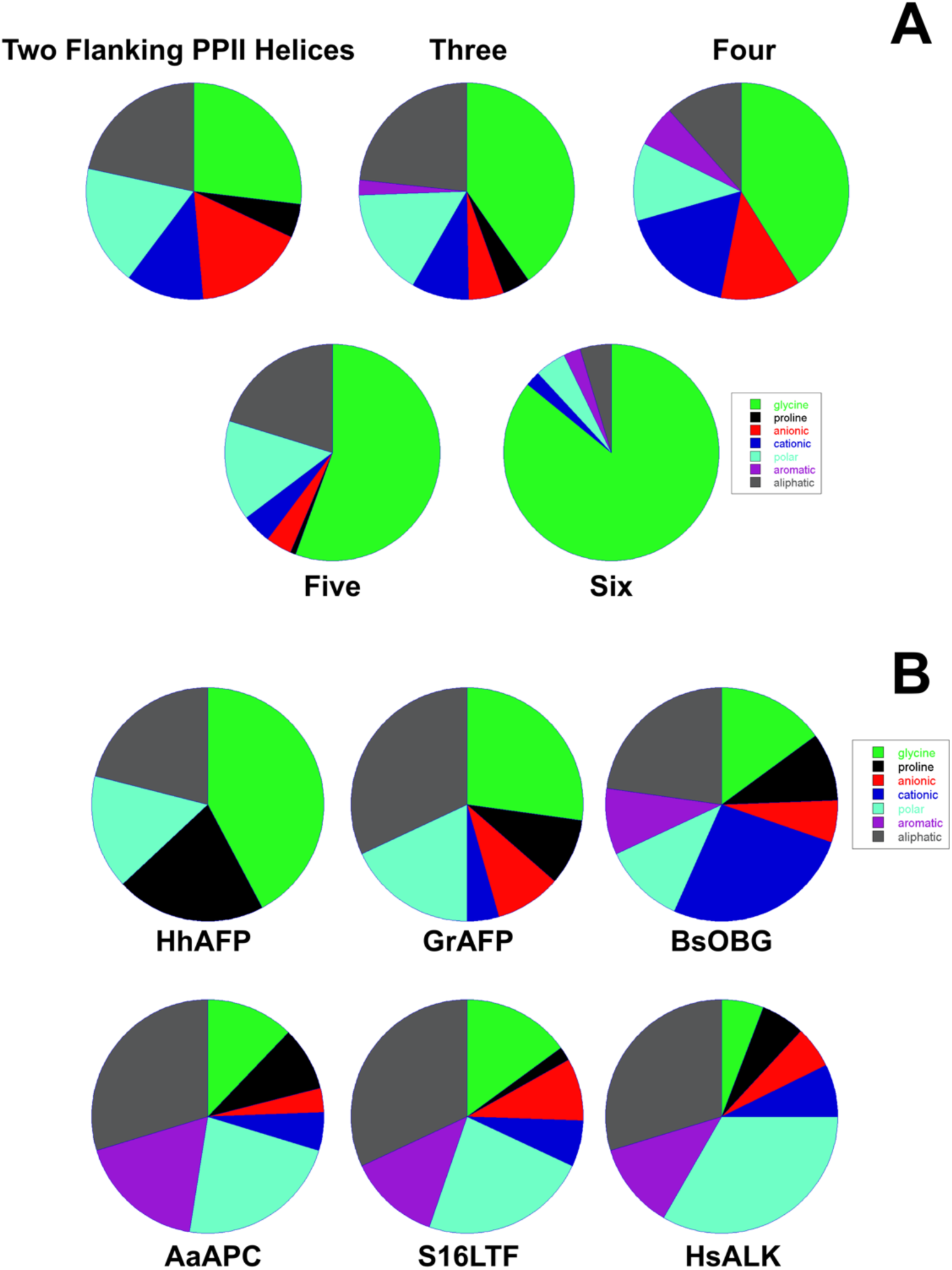
Glycine Content Increases with the Number of Neighboring PPII helices and Glycine-Rich PPII Helical Assemblies are Tolerant of Diverse Flanking Sequences. **A.** The Glycine content increases when a PPII helix is surrounded by more PPII helices. Residue color code: Gly=light green, Pro=black, anionic (Asp, Glu)=red, cationic (Arg, Lys, His)=blue, polar (Thr, Ser, Asn, Gln, Cys-SH)=cyan, aromatic (Phe, Tyr, Trp)=purple, aliphatic (Ala, Met, Val, Leu, Ile, Cys S-S)=gray. **B.** Glycine-rich PPII helical assemblies are robust enough to remain folded in the context of distinct flanking sequences, which can be rich in glycine and proline residues in the case of *Hh*AFP, cationic or aromatic residues for *Bs*OBG or *Aa*APC, respectively. The color code is the same as that used in panel **A**.

Human anaplastic lymphoma kinase is a receptor tyrosine kinase that plays key physiological roles in neuronal development and memory^72^ although its aberrant signaling is linked to obesity and cancer ^73^. The extracellular glycine-rich domain is key for binding peptide activators ^74^ and is composed of a remarkably large bundle of fourteen PPII helices ^64,65,75^. Three of these helices are surrounded by other PPII helices and are composed almost exclusively of glycine residues^64, 65,75^ (**Figure 7**). Some of the segments connecting the PPII helices are very short and consist of just a few residues Other connectors are long contain loops, β-hairpins or α-helices (**Figure 7**). Interestingly, the homologous protein in *C. elegans* (PDB: 7LIR) has ten somewhat shorter PPII helices arranged in a similar but not identical topology^75^.

The leukocyte tyrosine kinase (PDB: 7NX1)^65^ also contains a domain with fourteen glycine-rich polyproline II helices and its structure, variety of flanking sequences, preference for anti-parallel helix orientations and extensive H-bond network resemble those of the human anaplastic lymphoma kinase^64,65,75^. As previously proposed^65^, the complexity of this domain’s topology evinces its potential as a scaffold for protein design.

### Impact of flanking segments

Residues in flanking segments have well known to impact the formation and stability of β-sheet assemblies in amyloids ^68^. For example, Ala2 is on the edge of the Aβ amyloid structure and undergoes transitory unfolding events^76^. Nevertheless, substitution of Ala2 by Val is impactful, being protective against Alzheimer’s disease when heterozygous and disease-linked when homozygous ^77^ while A2T is protective ^78,79^. In the case of the short *Hh*AFP, all flanking regions and loops connecting the PPII helices are short (2 – 4 residues only) and rich in small polar residues; namely Gly, Cys, Asn and Thr, as well as Pro, which is a good turn forming residue (**Figure 2**, **Figure 8B**). This same trend is seen in a homologous six PPII helix AFP from *Megaphorura arctica*^80^ as well as in the longer antifreeze protein *Gr*AFP^57^, although it also contains three Val and one Leu residue in the connecting turns. Substitution of these highly exposed hydrophobic residues by polar residues to remove a reserve hydrophobic effect ^81^ is expected to be stabilizing and could be tested experimentally. In the case of *Aa*Apc, the content of aromatic flanking residues is notable and is consistent with this PPII helical assembly being mostly buried inside the protein. Interestingly, one of the aromatic flanking residues (W440) contacts a buried potassium ion forming a cation-π interaction. Some of the PPII helices; namely helix 7 of GsAFP, helix 4 of AaAPC, helices 3 and 4 of S16LTF, and helix 1 of *Hs*ALK have a cationic residue right at the C-terminus or in the first two residues following it. These residues are expected to contribute favorable charge-PPII helix macrodipole interactions^82^. With regard to the PPII domain of Obg, which is a translation regulator, its flanking residues are notably rich in cationic residues and play key roles in recognizing the ribosomal peptidyl transferase center ^83^.

### Packing Density

The formation of hydrogen bonds by buried peptide groups was reported to increase their packing density^84^. This strengthens van der Waals interactions and contributes favorably to protein conformational stability^83^. The short NH··OC hydrogen bond distances in *Hh*AFP’s polyproline II helices compared to those of α-helices and β-sheets reported previously^66^ led us to consider whether its packing density is higher. Interestingly, the *Hh*AFP does have a high packing density relative to other glycine rich PPII helical bundle proteins (**Table 2**). Nevertheless, a comparison of packing densities suggests that in general, glycine rich PPII helical bundles have a packing density similar to proteins composed of α-helices and β-sheet secondary structures (**Table 2**). This would imply that energetic contributions from van der Waals interactions would be similar.

**Table 2:** Protein Packing Densities

| **Protein** | **PDB File** | **Secondary Structure** | **Packing Density (Liang & Dill, 2001)** | **Packing Density (Measured here with Voronoia)** |
| --- | --- | --- | --- | --- |
| Myoglobin | 1MBO | α-helices | 0.716 | 0.729 |
| Lysozyme | 1LZL | α-helices (mostly) + some β-strands | 0.725 | 0.719 |
| Ribonuclease A | 7RSA | Mostly β-strands, three α-helices | 0.725 | 0.791 |
| SH3 domain | 1PGX | Mostly β-strands, one α-helix | 0.762 | 0.740 |
| *AaAFP* | 3BOB | PPII helical bundle |  | 0.751 |
| *GrAFP* | 7JJV | PPII helical bundle |  | 0.743 |
| *BsObg* | 1LNZ | PPII helical bundle |  | 0.742 |
| *TtObg* | 1UDX | PPII helical bundle |  | 0.738 |
| *EcObg* | 5M0Y | PPII helical bundle |  | 0.728 |
| *AaApc* | 5L9W | PPII helical bundle |  | 0.705 |
| *XaApc* | 5SVC | PPII helical bundle |  | 0.707 |
| S16LTF | 6F45 | PPII helical bundle |  | 0.732 |
| *HsALK* | 7N00 | PPII helical bundle |  | 0.720 |

### Parallel versus Antiparallel Configuration

Polyproline II helices have a macrodipole with an excess of positive charge near the N-terminus and negative charge near the C-terminus ^85^. A similar macrodipole is present in α-helices and a strong tendency towards antiparallel arrangements has been observed in proteins composed of four long α-helices arranged in a bundle^86^. This led us to assess the PPII helix orientation in glycine-rich PPII helical bundle domains. We found that there is a clear trend towards antiparallel orientation as 96% of successive PPII helices have their N-terminus placed on the same side as the C-terminus of the preceding PPII helix (**Figure 2-7**). Indeed, no PPII helix in this set of proteins has all parallel neighbors, all have at least one antiparallel neighbor, and all PPII helices have at least half of their neighbors in an antiparallel orientation (**Sup. Table 1**). Antiparallel orientations place helix macrodipoles in energetically favorable configurations and could thereby contribute to the conformational stability of these proteins.

### Putative Networks of C*α*-H*···*O=C H-bonds

Using the geometry criteria (<u>Cα-</u> <u>H··O</u>=C distance and bond angle cutoffs of 3.75 Å and 135°) described in the Methods Section, we identified 44 potential Cα-H···O=C H-bonds in *Gs*AFP, 15 in *Bs*Obg, 10 in *Ac*APC, 26 in S16LTF and 37 in *Hs*ALK. A complete list of the donors, acceptors, distances and angles of these H-bonds is listed in **Sup. Table 2**. The somewhat lower number of Cα-H···O=C H-bonds are present in *Aa*APC (**Sup. Table 2**) may be due to the irregular geometry of helices 1 and 6. Moreover, the glycine-rich PPII helical bundle domain of AaAPC is mostly buried. Thus, it may be stabilized by contacts with the surrounding protein elements and therefore require fewer hydrogen bonds. Whereas the existence of these H-bonds should be corroborated by other methods such as NMR spectroscopy in the future, and even though these H-bonds are weaker than canonical N-H··O=C H-bonds, due to their large numbers these Cα-H···O=C H-bonds are likely to provide an important source of conformational stability.

## Conclusions

In the last twenty years, the 3D structures of several proteins or protein domains formed by glycine-rich polyproline II helical bundles have been determined. By comparing and analyzing these structures, we have found that they share some common architectonic principles; in particular, they: 1) are tolerant of different loop lengths and flanking residues, 2) are as tightly packed as proteins composed of α-helices and β-sheets, 3) strongly prefer anti-parallel configurations and 4) generally have extensive networks of Cα-H···O=C hydrogen bonds. Due to the three dimensional and inter-helical nature of their NH··O=C and Cα-H···O=C hydrogen bonds, glycine-rich polyproline II domains have an internal stiffness and resistance to twisting or bending that sets them apart from α-helices and β-sheets and could make them a novel structural element for protein design. Protein designers have already started to incorporate polyproline II helices as a building block ^87^ and a binding target ^88^, but have not grafted a glycine-rich polyproline II helical bundle domain yet. Future work in this direction could well benefit from a consideration of the architectonic principles identified here.

## Author Contributions

**C. Segura Rodríguez:** Investigation, Data Curation, Validation, Writing-Original Draft Preparation, Writing-Reviewing and Editing. **D. V. Laurents**: Conceptualization, Funding Acquisition, Project Administration, Investigation, Data Curation, Validation, Writing-Original Draft Preparation, Writing-Reviewing and Editing.

## Dedication

This article is dedicated to Prof. Michael Levitt, an outstanding teacher and mentor and the inspriation of this study.

## Acknowledgement

This study is part of the project PID2022-137806OB-I00, funded by the Spanish Ministry of Science, Innovation and Universities: MICIN/AEI/10.13039/501100011033/FEDER, UE. The funding agency had no role in the study design or the interpretation of the data.

---

**Supporting Table 1:**
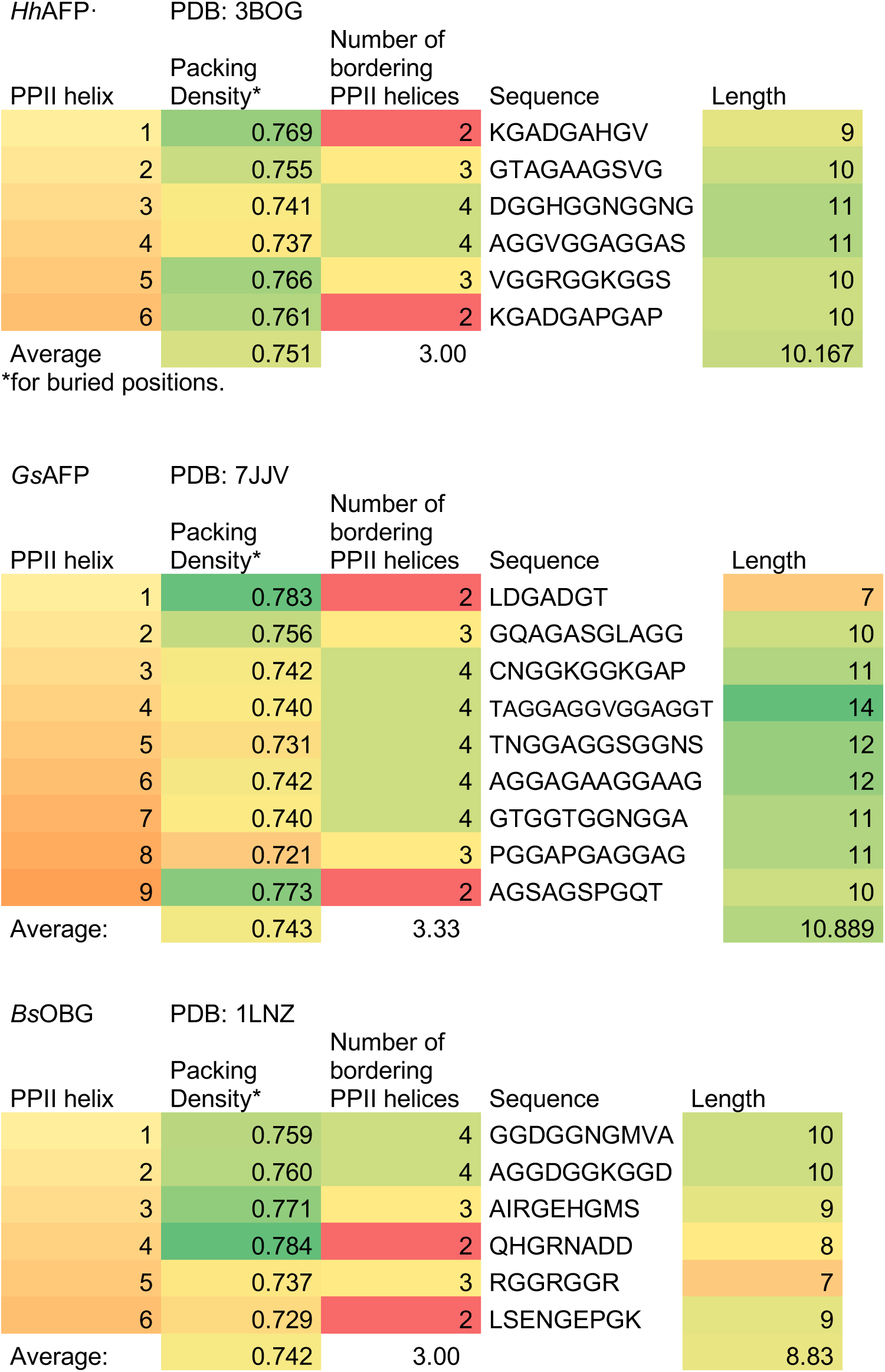
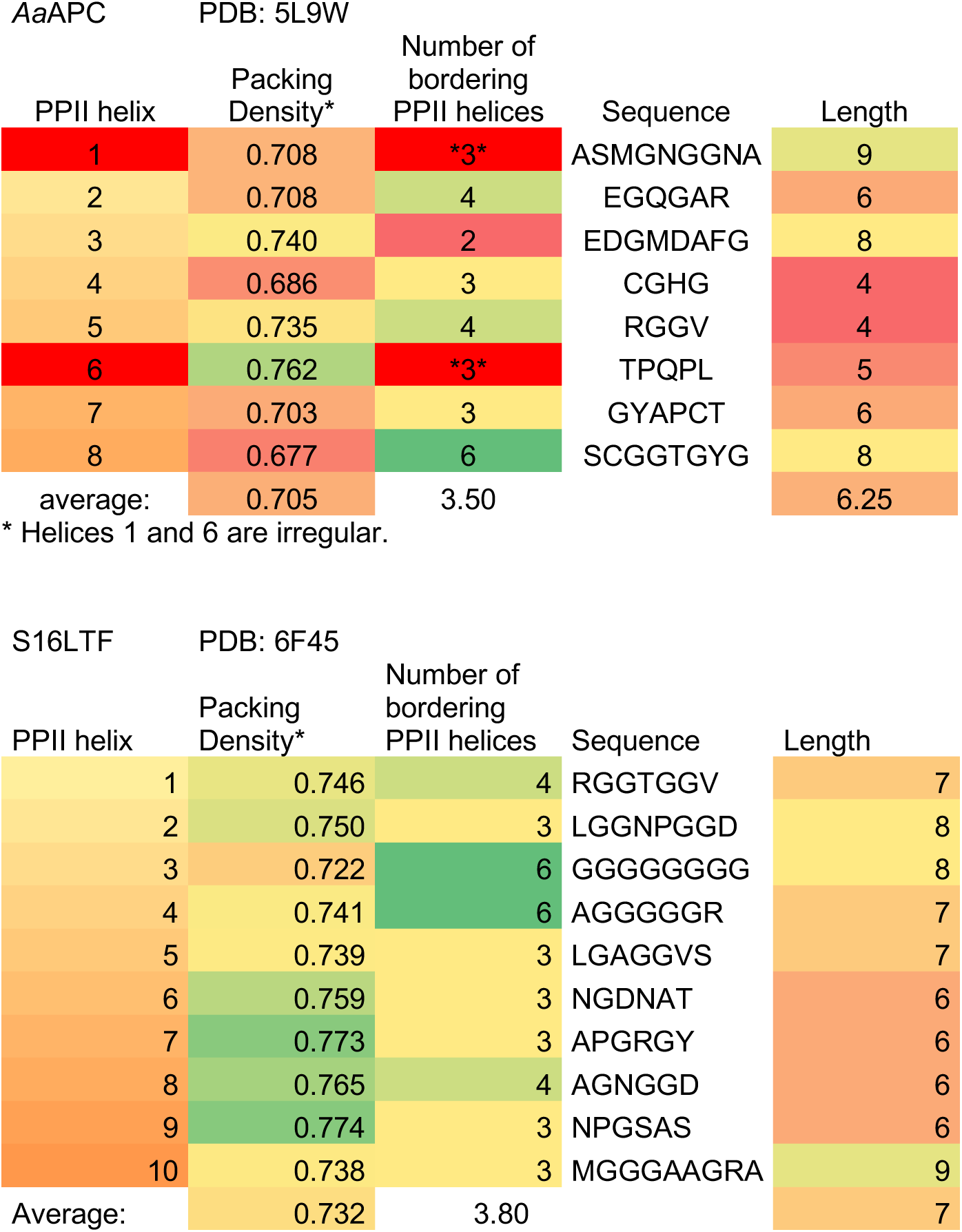
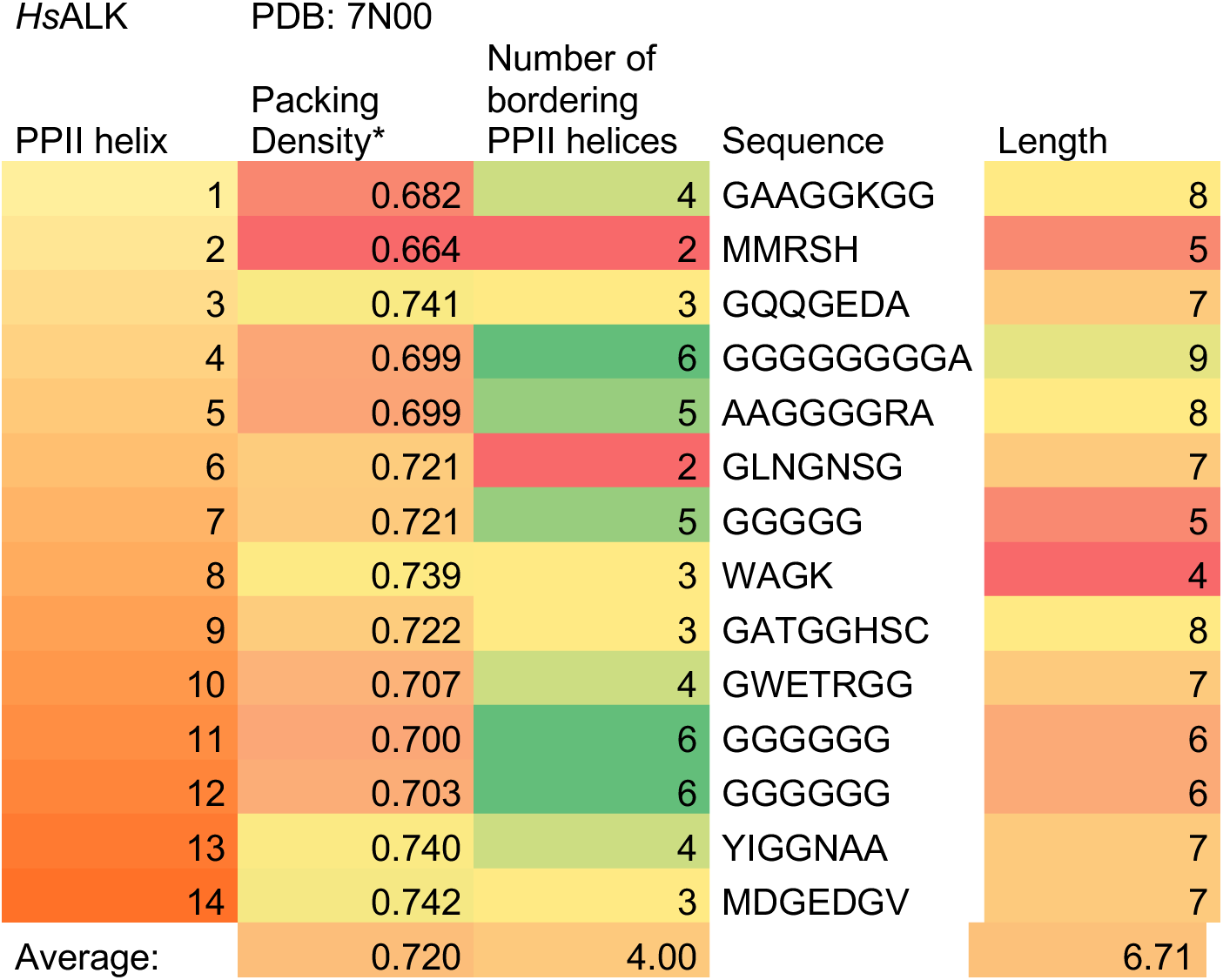
Packing Density of Buried PPII Helices

**Supporting Table 2:**
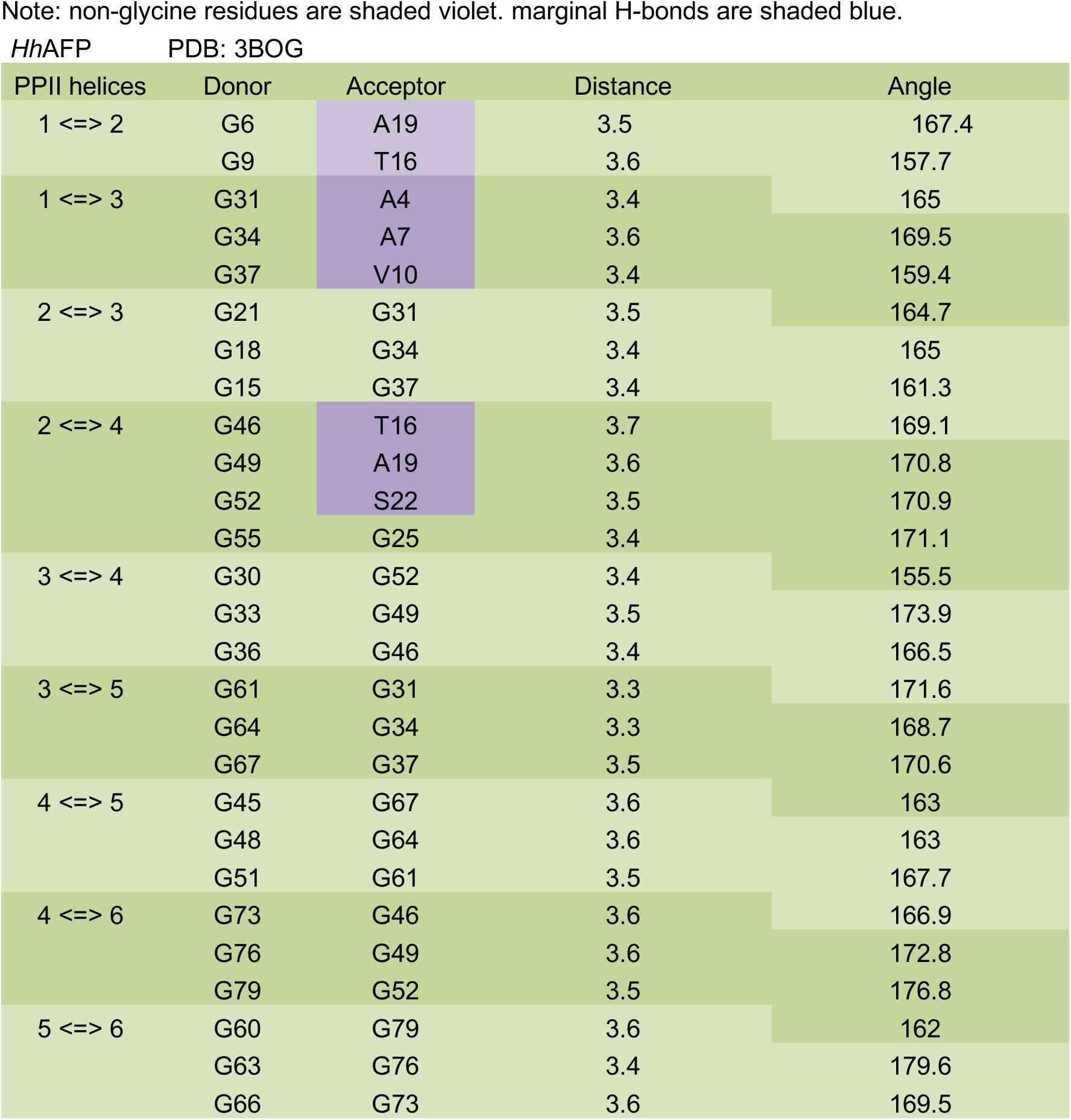
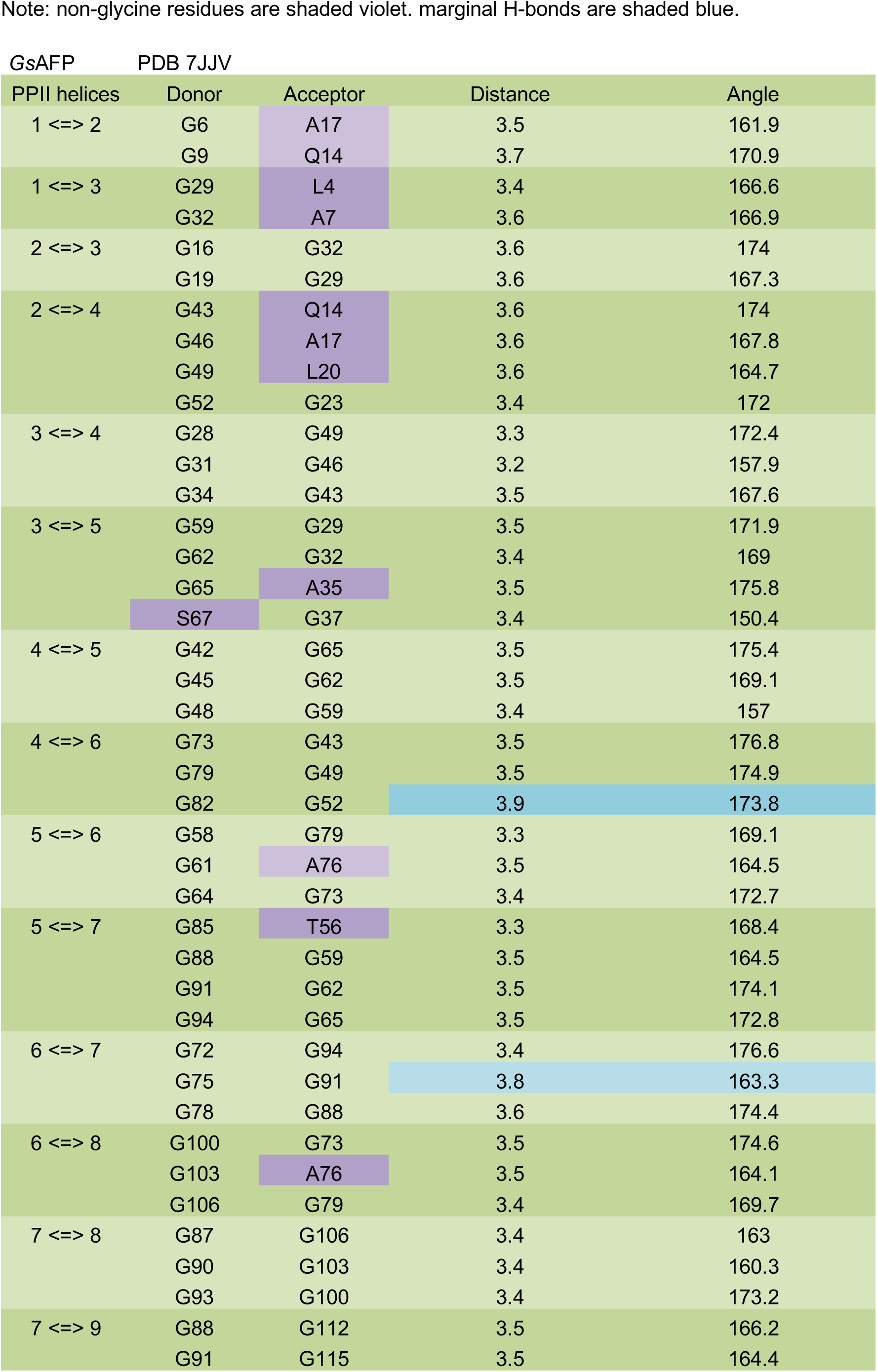
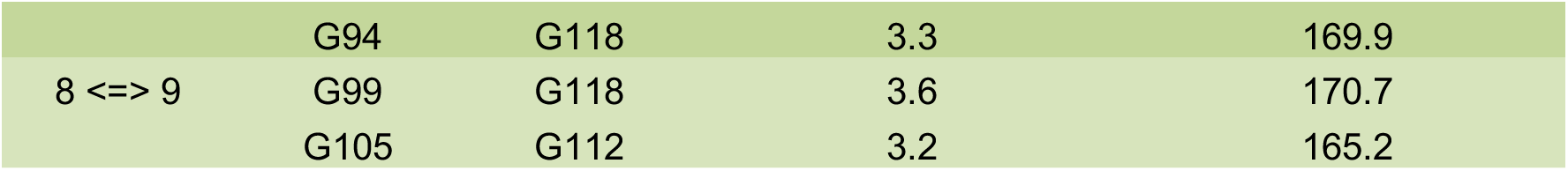
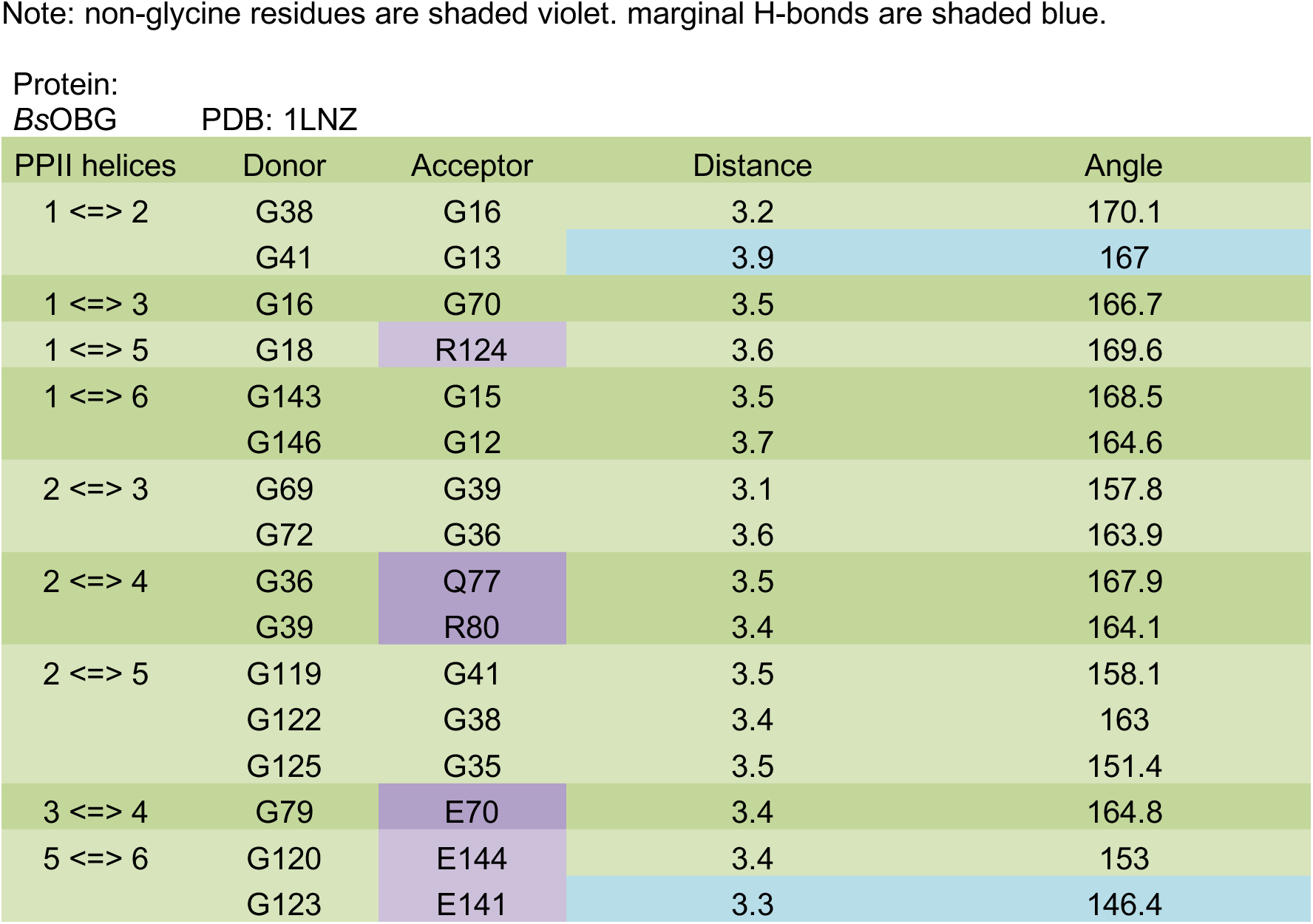
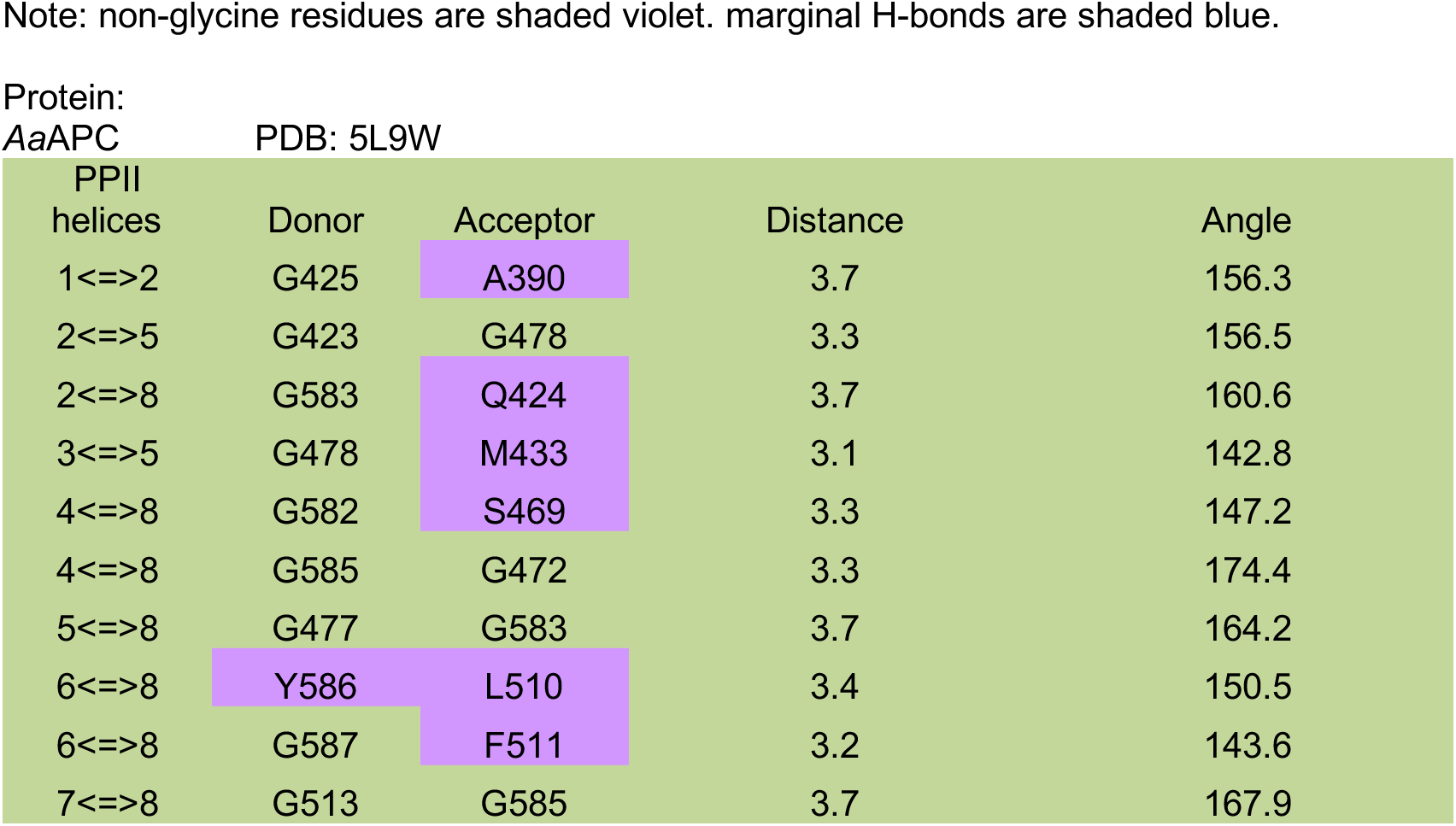
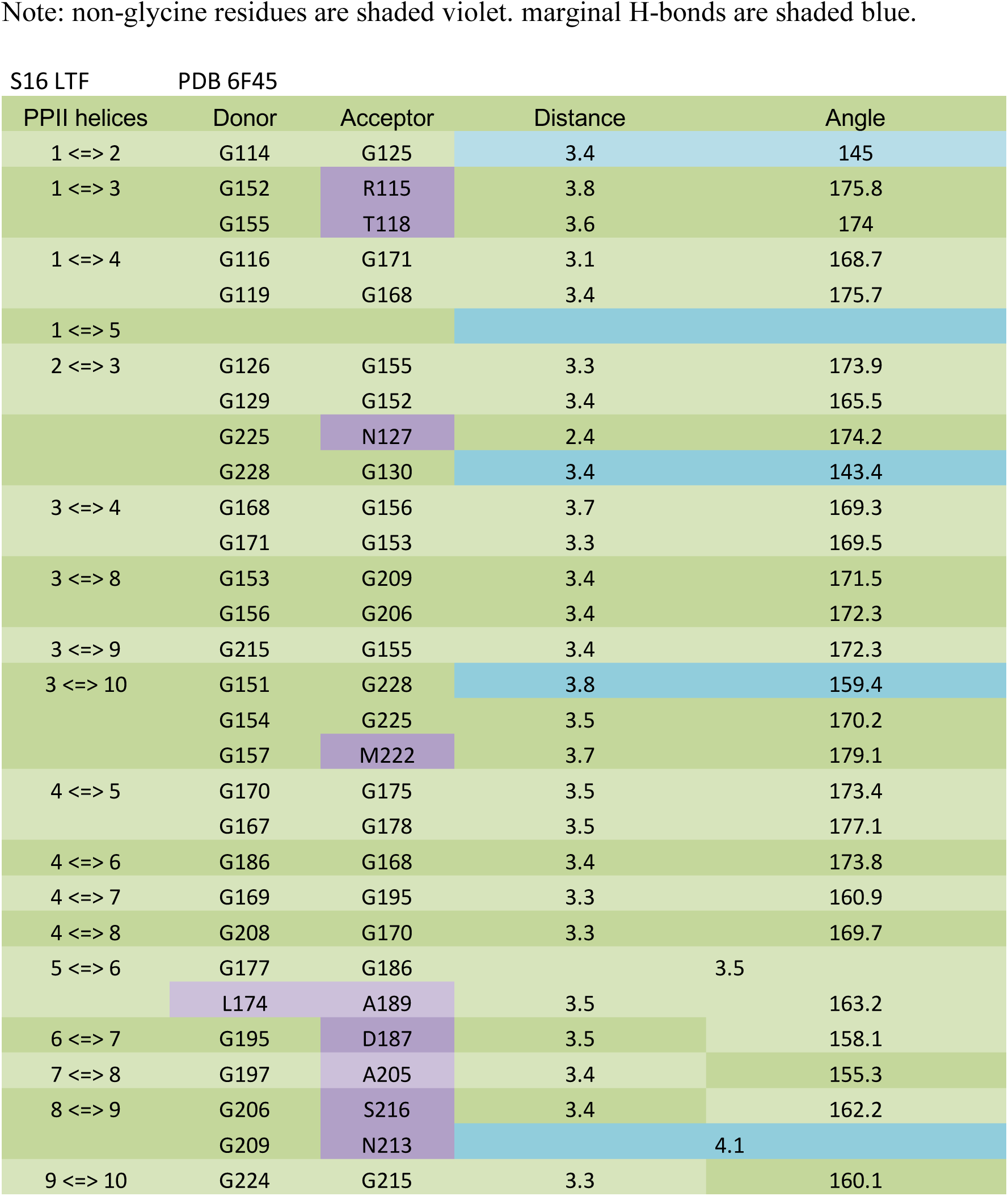
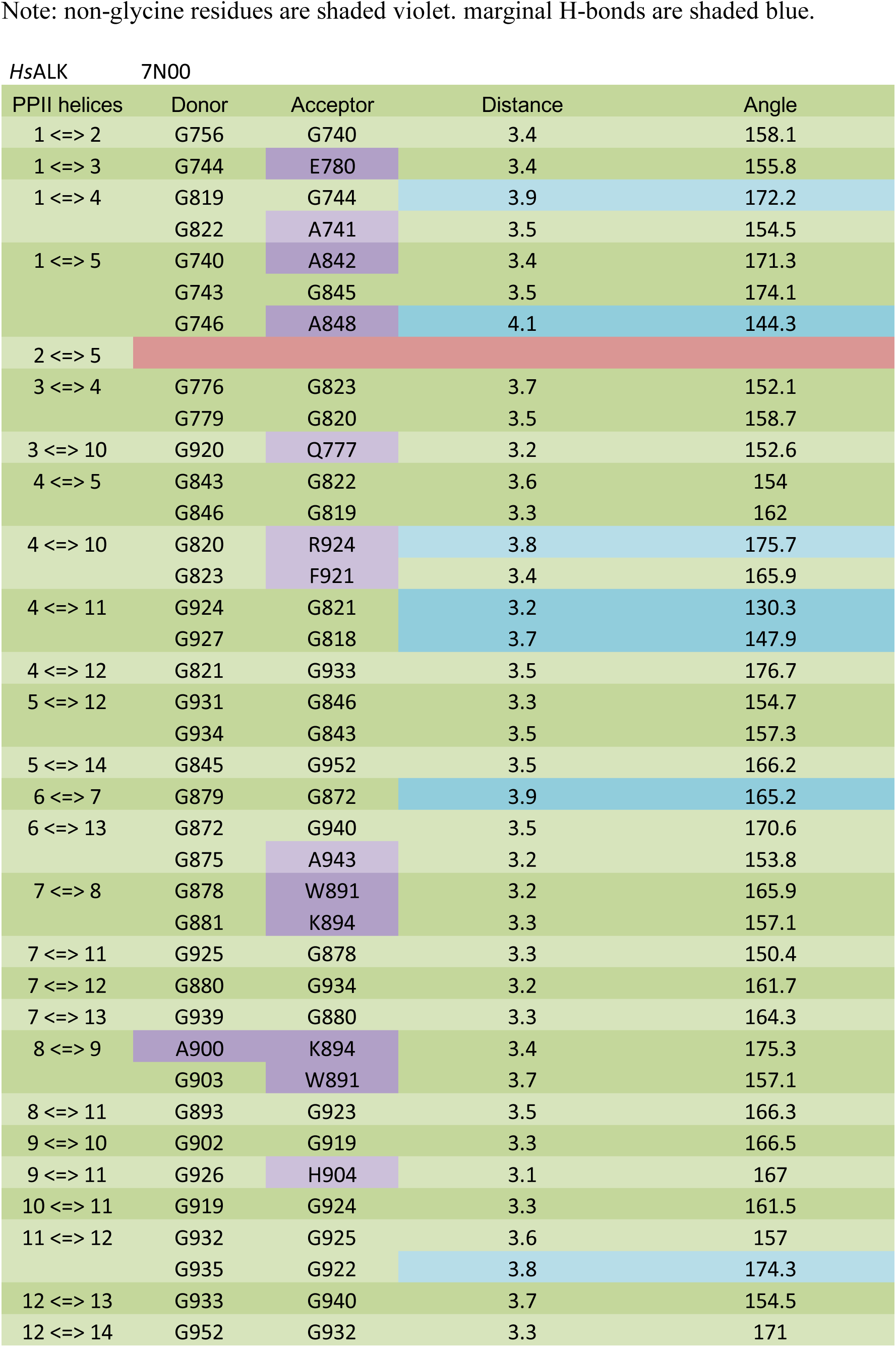
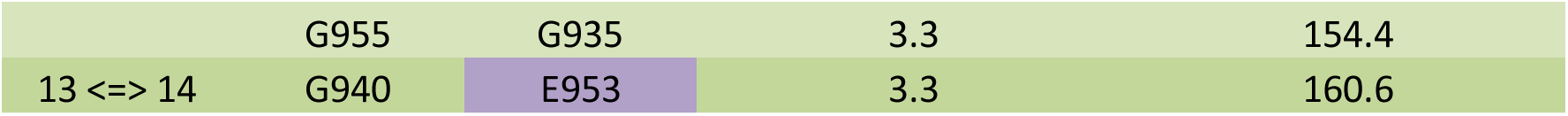
Cα-H···O=C Hydrogen Bonds

| PPII helices | Donor | Acceptor | Distance | Angle |
| --- | --- | --- | --- | --- |
| 1 <=> 2 | G6 | A19 | 3.5 | 167.4 |
|  | G9 | T16 | 3.6 | 157.7 |
| 1 <=> 3 | G31 | A4 | 3.4 | 165 |
|  | G34 | A7 | 3.6 | 169.5 |
|  | G37 | V10 | 3.4 | 159.4 |
| 2 <=> 3 | G21 | G31 | 3.5 | 164.7 |
|  | G18 | G34 | 3.4 | 165 |
|  | G15 | G37 | 3.4 | 161.3 |
| 2 <=> 4 | G46 | T16 | 3.7 | 169.1 |
|  | G49 | A19 | 3.6 | 170.8 |
|  | G52 | S22 | 3.5 | 170.9 |
|  | G55 | G25 | 3.4 | 171.1 |
| 3 <=> 4 | G30 | G52 | 3.4 | 155.5 |
|  | G33 | G49 | 3.5 | 173.9 |
|  | G36 | G46 | 3.4 | 166.5 |
| 3 <=> 5 | G61 | G31 | 3.3 | 171.6 |
|  | G64 | G34 | 3.3 | 168.7 |
|  | G67 | G37 | 3.5 | 170.6 |
| 4 <=> 5 | G45 | G67 | 3.6 | 163 |
|  | G48 | G64 | 3.6 | 163 |
|  | G51 | G61 | 3.5 | 167.7 |
| 4 <=> 6 | G73 | G46 | 3.6 | 166.9 |
|  | G76 | G49 | 3.6 | 172.8 |
|  | G79 | G52 | 3.5 | 176.8 |
| 5 <=> 6 | G60 | G79 | 3.6 | 162 |
|  | G63 | G76 | 3.4 | 179.6 |
|  | G66 | G73 | 3.6 | 169.5 |

| GsAFP | PDB 7JJV |  |  |  |
| --- | --- | --- | --- | --- |
| PPII helices | Donor | Acceptor | Distance | Angle |
| 1 <=> 2 | G6 | A17 | 3.5 | 161.9 |
|  | G9 | Q14 | 3.7 | 170.9 |
| 1 <=> 3 | G29 | L4 | 3.4 | 166.6 |
|  | G32 | A7 | 3.6 | 166.9 |
| 2 <=> 3 | G16 | G32 | 3.6 | 174 |
|  | G19 | G29 | 3.6 | 167.3 |
| 2 <=> 4 | G43 | Q14 | 3.6 | 174 |
|  | G46 | A17 | 3.6 | 167.8 |
|  | G49 | L20 | 3.6 | 164.7 |
|  | G52 | G23 | 3.4 | 172 |
| 3 <=> 4 | G28 | G49 | 3.3 | 172.4 |
|  | G31 | G46 | 3.2 | 157.9 |
|  | G34 | G43 | 3.5 | 167.6 |
| 3 <=> 5 | G59 | G29 | 3.5 | 171.9 |
|  | G62 | G32 | 3.4 | 169 |
|  | G65 | A35 | 3.5 | 175.8 |
|  | S67 | G37 | 3.4 | 150.4 |
| 4 <=> 5 | G42 | G65 | 3.5 | 175.4 |
|  | G45 | G62 | 3.5 | 169.1 |
|  | G48 | G59 | 3.4 | 157 |
| 4 <=> 6 | G73 | G43 | 3.5 | 176.8 |
|  | G79 | G49 | 3.5 | 174.9 |
|  | G82 | G52 | 3.9 | 173.8 |
| 5 <=> 6 | G58 | G79 | 3.3 | 169.1 |
|  | G61 | A76 | 3.5 | 164.5 |
|  | G64 | G73 | 3.4 | 172.7 |
| 5 <=> 7 | G85 | T56 | 3.3 | 168.4 |
|  | G88 | G59 | 3.5 | 164.5 |
|  | G91 | G62 | 3.5 | 174.1 |
|  | G94 | G65 | 3.5 | 172.8 |
| 6 <=> 7 | G72 | G94 | 3.4 | 176.6 |
|  | G75 | G91 | 3.8 | 163.3 |
|  | G78 | G88 | 3.6 | 174.4 |
| 6 <=> 8 | G100 | G73 | 3.5 | 174.6 |
|  | G103 | A76 | 3.5 | 164.1 |
|  | G106 | G79 | 3.4 | 169.7 |
| 7 <=> 8 | G87 | G106 | 3.4 | 163 |
|  | G90 | G103 | 3.4 | 160.3 |
|  | G93 | G100 | 3.4 | 173.2 |
| 7 <=> 9 | G88 | G112 | 3.5 | 166.2 |
|  | G91 | G115 | 3.5 | 164.4 |
|  | G94 | G118 | 3.3 | 169.9 |
| 8 <=> 9 | G99 | G118 | 3.6 | 170.7 |
|  | G105 | G112 | 3.2 | 165.2 |
| 1 <=> 2 | G38 | G16 | 3.2 | 170.1 |
|  | G41 | G13 | 3.9 | 167 |
| 1 <=> 3 | G16 | G70 | 3.5 | 166.7 |
| 1 <=> 5 | G18 | R124 | 3.6 | 169.6 |
| 1 <=> 6 | G143 | G15 | 3.5 | 168.5 |
|  | G146 | G12 | 3.7 | 164.6 |
| 2 <=> 3 | G69 | G39 | 3.1 | 157.8 |
|  | G72 | G36 | 3.6 | 163.9 |
| 2 <=> 4 | G36 | Q77 | 3.5 | 167.9 |
|  | G39 | R80 | 3.4 | 164.1 |
| 2 <=> 5 | G119 | G41 | 3.5 | 158.1 |
|  | G122 | G38 | 3.4 | 163 |
|  | G125 | G35 | 3.5 | 151.4 |
| 3 <=> 4 | G79 | E70 | 3.4 | 164.8 |
| 5 <=> 6 | G120 | E144 | 3.4 | 153 |
|  | G123 | E141 | 3.3 | 146.4 |

| PPII<br>helices | Donor | Acceptor | Distance | Angle |
| --- | --- | --- | --- | --- |
| 1<=>2 | G425 | A390 | 3.7 | 156.3 |
| 2<=>5 | G423 | G478 | 3.3 | 156.5 |
| 2<=>8 | G583 | Q424 | 3.7 | 160.6 |
| 3<=>5 | G478 | M433 | 3.1 | 142.8 |
| 4<=>8 | G582 | S469 | 3.3 | 147.2 |
| 4<=>8 | G585 | G472 | 3.3 | 174.4 |
| 5<=>8 | G477 | G583 | 3.7 | 164.2 |
| 6<=>8 | Y586 | L510 | 3.4 | 150.5 |
| 6<=>8 | G587 | F511 | 3.2 | 143.6 |
| 7<=>8 | G513 | G585 | 3.7 | 167.9 |

| S16 LTF | PDB 6F45 |  |  |  |
| --- | --- | --- | --- | --- |
| PPII helices | Donor | Acceptor | Distance | Angle |
| 1 <=> 2 | G114 | G125 | 3.4 | 145 |
| 1 <=> 3 | G152 | R115 | 3.8 | 175.8 |
|  | G155 | T118 | 3.6 | 174 |
| 1 <=> 4 | G116 | G171 | 3.1 | 168.7 |
|  | G119 | G168 | 3.4 | 175.7 |
| 1 <=> 5 |  |  |  |  |
| 2 <=> 3 | G126 | G155 | 3.3 | 173.9 |
|  | G129 | G152 | 3.4 | 165.5 |
|  | G225 | N127 | 2.4 | 174.2 |
|  | G228 | G130 | 3.4 | 143.4 |
| 3 <=> 4 | G168 | G156 | 3.7 | 169.3 |
|  | G171 | G153 | 3.3 | 169.5 |
| 3 <=> 8 | G153 | G209 | 3.4 | 171.5 |
|  | G156 | G206 | 3.4 | 172.3 |
| 3 <=> 9 | G215 | G155 | 3.4 | 172.3 |
| 3 <=> 10 | G151 | G228 | 3.8 | 159.4 |
|  | G154 | G225 | 3.5 | 170.2 |
|  | G157 | M222 | 3.7 | 179.1 |
| 4 <=> 5 | G170 | G175 | 3.5 | 173.4 |
|  | G167 | G178 | 3.5 | 177.1 |
| 4 <=> 6 | G186 | G168 | 3.4 | 173.8 |
| 4 <=> 7 | G169 | G195 | 3.3 | 160.9 |
| 4 <=> 8 | G208 | G170 | 3.3 | 169.7 |
| 5 <=> 6 | G177 | G186 |  | 3.5 |
|  | L174 | A189 | 3.5 | 163.2 |
| 6 <=> 7 | G195 | D187 | 3.5 | 158.1 |
| 7 <=> 8 | G197 | A205 | 3.4 | 155.3 |
| 8 <=> 9 | G206 | S216 | 3.4 | 162.2 |
|  | G209 | N213 |  | 4.1 |
| 9 <=> 10 | G224 | G215 | 3.3 | 160.1 |

| HsALK | 7N00 |  |  |  |
| --- | --- | --- | --- | --- |
| PPII helices | Donor | Acceptor | Distance | Angle |
| 1 <=> 2 | G756 | G740 | 3.4 | 158.1 |
| 1 <=> 3 | G744 | E780 | 3.4 | 155.8 |
| 1 <=> 4 | G819 | G744 | 3.9 | 172.2 |
|  | G822 | A741 | 3.5 | 154.5 |
| 1 <=> 5 | G740 | A842 | 3.4 | 171.3 |
|  | G743 | G845 | 3.5 | 174.1 |
|  | G746 | A848 | 4.1 | 144.3 |
| 2 <=> 5 |  |  |  |  |
| 3 <=> 4 | G776 | G823 | 3.7 | 152.1 |
|  | G779 | G820 | 3.5 | 158.7 |
| 3 <=> 10 | G920 | Q777 | 3.2 | 152.6 |
| 4 <=> 5 | G843 | G822 | 3.6 | 154 |
|  | G846 | G819 | 3.3 | 162 |
| 4 <=> 10 | G820 | R924 | 3.8 | 175.7 |
|  | G823 | F921 | 3.4 | 165.9 |
| 4 <=> 11 | G924 | G821 | 3.2 | 130.3 |
|  | G927 | G818 | 3.7 | 147.9 |
| 4 <=> 12 | G821 | G933 | 3.5 | 176.7 |
| 5 <=> 12 | G931 | G846 | 3.3 | 154.7 |
|  | G934 | G843 | 3.5 | 157.3 |
| 5 <=> 14 | G845 | G952 | 3.5 | 166.2 |
| 6 <=> 7 | G879 | G872 | 3.9 | 165.2 |
| 6 <=> 13 | G872 | G940 | 3.5 | 170.6 |
|  | G875 | A943 | 3.2 | 153.8 |
| 7 <=> 8 | G878 | W891 | 3.2 | 165.9 |
|  | G881 | K894 | 3.3 | 157.1 |
| 7 <=> 11 | G925 | G878 | 3.3 | 150.4 |
| 7 <=> 12 | G880 | G934 | 3.2 | 161.7 |
| 7 <=> 13 | G939 | G880 | 3.3 | 164.3 |
| 8 <=> 9 | A900 | K894 | 3.4 | 175.3 |
|  | G903 | W891 | 3.7 | 157.1 |
| 8 <=> 11 | G893 | G923 | 3.5 | 166.3 |
| 9 <=> 10 | G902 | G919 | 3.3 | 166.5 |
| 9 <=> 11 | G926 | H904 | 3.1 | 167 |
| 10 <=> 11 | G919 | G924 | 3.3 | 161.5 |
| 11 <=> 12 | G932 | G925 | 3.6 | 157 |
|  | G935 | G922 | 3.8 | 174.3 |
| 12 <=> 13 | G933 | G940 | 3.7 | 154.5 |
| 12 <=> 14 | G952 | G932 | 3.3 | 171 |
|  | G955 | G935 | 3.3 | 154.4 |
| 13 <=> 14 | G940 | E953 | 3.3 | 160.6 |

